# Data-driven denoising in spinal cord fMRI with principal component analysis

**DOI:** 10.1101/2025.01.23.634596

**Authors:** Kimberly J. Hemmerling, Andrew D. Vigotsky, Charlotte Glanville, Robert L. Barry, Molly G. Bright

## Abstract

Numerous approaches have been used to denoise spinal cord functional magnetic resonance imaging (fMRI) data. Principal component analysis (PCA)-based techniques, which derive regressors from a noise region of interest (ROI), have been used in both brain (e.g., CompCor) and spinal cord fMRI. However, spinal cord fMRI denoising methods have yet to be systematically evaluated. Here, we formalize and evaluate a PCA-based technique for deriving nuisance regressors for spinal cord fMRI analysis (SpinalCompCor). In this method, regressors are derived with PCA from a noise ROI, an area defined outside of the spinal cord and cerebrospinal fluid. A parallel analysis is used to systematically determine how many components to retain as regressors for modeling; this designated a median of 9 regressors across four fMRI datasets: motor task (n=26), breathing task (n=27), and resting state (n=15 and n=10). First-level fMRI modeling demonstrated that principal component regressors did fit noise (e.g., physiological noise from blood vessels), though the effectiveness may be dependent upon the acquisition parameters. However, group-level activation maps did not show a clear benefit from including SpinalCompCor regressors. The potential for collinearity of principal component regressors with the task may be a concern, and this should be considered in future implementations for which task-correlated noise is anticipated. In general, denoising with SpinalCompCor regressors in place of physiological recording-derived regressors is only recommended when the latter are unavailable, as SpinalCompCor may not consistently reproduce recording-based denoising across datasets or acquisitions.

## 1 INTRODUCTION

Spinal cord functional magnetic resonance imaging (fMRI) can provide valuable insight into neural processes of the human spinal cord; however, well-known challenges include the influence of physiological noise and motion. Spinal cord voxels are highly susceptible to artifacts from cardiac and respiratory sources (Eippert et al., 2017). The nearby dilation of arteries after systole as well as the cardiac-linked pulsatile flow of cerebrospinal fluid (CSF) surrounding the cord confound the fMRI signal timeseries (Piché et al., 2009). Additionally, nearby inflation of the lungs and movement of the chest cavity during ventilation greatly impacts the homogeneity of the B_0_ magnetic field and, thus, fMRI signals within the spinal cord itself (Raj et al., 2001).

Robust approaches to denoising are of utmost importance for spinal cord fMRI. One common technique for combatting physiological noise confounds is RETROICOR, a physiological recording-based denoising technique developed for brain fMRI (Glover et al., 2000), with adaptations to the spinal cord (Brooks et al., 2008; Kong et al., 2012). Fourier-based RETROICOR regressors are modeled from cardiac and respiratory signals externally recorded during scanning and then regressed out of the fMRI data. Data-driven nuisance regressors can also be generated in specific regions of interest (ROIs) (e.g., CSF (Brooks et al., 2008; Kong et al., 2012)) or via independent component analysis (ICA) methods (Hu et al., 2018; Xie et al., 2012). Motion correction and the subsequent use of motion regressors in fMRI models are commonplace (see Eippert et al. 2017 for a detailed review of approaches to denoising in spinal cord fMRI (Eippert et al., 2017)).

Another approach is to use principal component analysis (PCA) within a subset of imaging voxels to derive multiple nuisance regressors for fMRI denoising. CompCor, a PCA-based denoising technique for brain fMRI, defines a noise ROI in which PCA is applied to obtain nuisance regressors (Behzadi et al., 2007). This noise ROI is defined either anatomically (i.e., in white matter and CSF) or based on voxel timeseries with the highest temporal standard deviation. Another method uses PCA in brain edge voxels to define regressors and improve removal of motion confounds in resting state brain fMRI data (Patriat et al., 2015). This type of data-driven PCA-based approach for denoising can be advantageous when physiological recordings are low quality or unavailable, such as in clinical studies.

While ICA-based methods are popular in fMRI analyses, PCA-based methods remain a common choice for defining regressors. (See Caballero-Gaudes and Reynolds for a detailed discussion of ICA and PCA denoising methods (Caballero-Gaudes & Reynolds, 2017).) The current work focuses on PCA as a denoising strategy for a few primary reasons. First, principal components are inherently ordered, which facilitates an objective selection of regressors, for example, by defining criteria based on a cumulative variance threshold. This approach offers an advantage over some ICA-based techniques that can require more complex or labor-intensive component classification methods. Also, PCA requires less computational power than ICA.

CompCor methods for brain fMRI data have inspired related methods for spinal cord fMRI analysis. For example, in an implementation of PCA-based denoising for 7 T resting state spinal cord fMRI, a noise ROI was defined as anywhere external to the cord and its adjacent CSF; then, PCA of voxel timeseries within this noise ROI was used to produce nuisance regressors (Barry et al., 2014, 2016). In this technique, components were selected based on the percent of signal variance explained and were then regressed out from the data prior to motion correction to improve its efficacy. Although this method has been previously applied, it has yet to be rigorously evaluated for spinal cord fMRI more generally.

In this work, we extend on the method initially proposed for 7 T resting state spinal cord fMRI (Barry et al., 2014, 2016), and we will henceforth refer to PCA-based denoising for spinal cord fMRI as “SpinalCompCor”. For SpinalCompCor denoising, we assume that signal outside of the cord is non-neural and that this noise similarly affects the fMRI timeseries both within the spinal cord and in surrounding tissues. Under this assumption, principal components can be derived from the timeseries in these typically unused voxels and harnessed as a tool for denoising of spinal cord fMRI data.

We first define our method to calculate the principal components from a noise ROI in spinal cord fMRI data. It is important to identify meaningful principal components to include as regressors in the fMRI noise model. Thus, we employ a parallel analysis to determine, in a given dataset, how many components should be retained as nuisance regressors. This parallel analysis systematically determines how many components to retain by comparing principal component eigenvalues from real data to those from surrogate data, essentially identifying the number of components that explain greater variance than the surrogate components do by chance. The parallel analysis is evaluated in two 3 T task fMRI datasets and two 3 T resting state fMRI datasets. Subject-level fMRI analysis is subsequently performed in these four datasets to assess the benefit of including the selected SpinalCompCor regressors in fMRI modeling. Finally, we evaluate the effect of including data-driven SpinalCompCor regressors for group-level activation and connectivity mapping.

## 2 METHODS

Four previously collected spinal cord fMRI datasets were used in this study, including a hand-grasping task, a breath-holding task, and two resting state acquisitions. The isometric unilateral hand-grasping task was expected to evoke motor activation localized to the ventral horns of the C7-C8 spinal cord segments within the field of view (Hemmerling et al., 2023a; Vanderah & Gould, 2016). The breath-holding task was designed to induce hypercapnia and evoke systemic vasodilation, from which spinal cord vascular reactivity could be estimated (Hemmerling et al., 2025). The resting state acquisitions were expected to be sensitive to intrinsic signal fluctuations and functional networks in the spinal cord (Barry et al., 2018; Weber et al., 2018). In all four datasets, we expect SpinalCompCor regressors to capture noise associated with physiology, non-physiological motion, and other artifacts. The hand-grasping and breath-holding tasks may additionally exhibit task-correlated motion as well as physiological changes (e.g., heart rate, chest position) related to the task, while the resting state may be expected to have more subtle motion and physiological fluctuations. Thus, we aim to explore any differences in the selection of principal component regressors and model performance with the new regressors across a few fMRI acquisitions.

### 2.1 Data collection

Data were collected in a studies approved by either the Northwestern University Institutional Review Board (hand-grasping task, breath-holding task; IRB Protocol Number: STU00209651). Data shared by collaborators were approved by either the Northwestern University Institutional Review Board or the Vanderbilt University Institutional Review Board (resting state datasets). Informed consent was obtained from all participants for being included in the studies.

#### 2.1.1 Hand-grasping task fMRI

The hand-grasping task dataset was collected at the Northwestern University Center for Translational Imaging on a 3 T Siemens Prisma MRI system (Siemens Healthcare, Erlangen, Germany) with a 64-channel head/neck coil and SatPad™ cervical collar from 30 healthy participants (25.9 ± 4.5 years, 19 F) (Hemmerling et al., 2023b, 2023a). Four subjects were excluded because of excessive image artifacts, an incidental finding, and technical issues (included sample: n=26, 25.8 ± 4.5 years, 17 F).

A high-resolution T2-weighted sagittal anatomical MRI was acquired (repetition time (TR)/echo time (TE) = 1500/135 ms, sagittal slice thickness = 0.8 mm, in-plane resolution 0.39 mm^2^, 64 slices, flip-angle (FA) = 140°, field-of-view (FOV) = 640 mm^2^). The functional sequence was a gradient-echo echo-planar imaging (EPI) sequence with ZOOMit selective excitation (TR/TE = 2000/30 ms, 1 × 1 × 3 mm^3^, 25 slices, FA = 90°, FOV = 128 × 44 mm^2^). Cervical cord coverage was approximately the C4-C7 vertebral levels.

The 10-minute (300 volume) hand-grasping task run consisted of 24 trials of 15-second pseudo-randomized left and right grasps. Participants viewed real-time force feedback while squeezing custom MRI-safe handgrips **(Fig. S1)**, targeting 25% maximum-voluntary contraction (MVC). Physiological data (respiration, pulse) were collected during the session. The first of the two fMRI runs acquired in the original study was used in the parallel analysis and subject-level modeling; both runs were used for group-level activation mapping. Respiration belt data could not be collected for one subject. This subject was excluded from fMRI modeling (see section 2.2.4) and following analyses (final included sample: n=25, 25.8 ± 4.6 years, 16 F).

#### 2.1.2 Breath-holding task fMRI

The breath-holding task dataset was collected from the same participants during the same sessions as the hand-grasping task (Hemmerling et al., 2024, 2025). Three subjects were excluded because of excessive image artifacts and an incidental finding (included sample: n=27, 25.7 ± 4.7 years, 18 F).

The 6-minute 50-second (205 volume) breath-holding task scan was acquired with the same parameters as the hand-grasping scan except for the number of volumes. The task run began with a 20-second rest, followed by 7 breath-holding cycles, and ended with a 30-second rest.

Each cycle had 24 seconds of paced breathing, an 18-second end-expiration breath-hold, a 2-second exhalation, and a 6-second recovery period. Physiological data (respiration, pulse, exhaled CO_2_) were collected during the session. The first of the two fMRI runs acquired in the original study was used here. Respiration belt data could not be collected for one subject. This subject was excluded from fMRI modeling (see section 2.2.4) and following analyses (final included sample: n=26, 25.7 ± 4.7 years, 17 F).

#### 2.1.3 Resting state fMRI

##### Resting State A

One resting state fMRI dataset was collected at the Northwestern University Center for Translational Imaging on a Siemens Prisma 3 T scanner with a 64-channel head/neck coil and SatPad™ cervical collar from 15 healthy participants (28.1 ± 2.4 years, 6 F) (Weber et al., 2016b, 2016a, 2018). A 6-minute 40-second (160 volume) gradient-echo EPI sequence with ZOOMit selective excitation was acquired (TR/TE = 2500/30 ms, 1 × 1 × 3 mm^3^, 31 slices, FA = 80°, FOV = 128 × 44 mm^2^). Cervical cord coverage was approximately from the C3 (inferior endplate) to T1 (superior endplate) vertebra. A high-resolution T2-weighted anatomical MRI was acquired using a single slab three-dimensional turbo spin echo sequence with a slab selective, variable excitation pulse (SPACE, TR/TE_eff_ = 1500/115 ms, echo train length = 78, FA=90°/140°, effective resolution 0.8 x 0.8 x 0.8 mm^3^, interpolated resolution 0.8 x 0.4 x 0.4 mm^3^. Physiological data (respiration, pulse) were collected during the session.

##### Resting State B

A second resting state dataset was collected at the Vanderbilt University Institute of Imaging Science on a Philips Achieva 3 T scanner (Best, The Netherlands) from 10 healthy participants (27.0 ± 4.9 years, 5 F) (Barry et al., 2018). A sagittal anatomical localizer scan was acquired. The functional sequence was a 3D gradient-echo multi-shot sequence (TR = 34 ms, TE = 8 ms, acquired voxel size = 1 × 1 × 5 mm^3^, interpolated voxel size 0.59 x 0.59 x 5 mm^3^, 12 slices, FA = 90°, FOV = 150 mm^2^, echo train length=7, SENSE = 2.0 (left-right), volume acquisition time (VAT) = 2.76 s). The cervical cord coverage was from approximately the C2-C5 vertebral level. A 20-minute (434 volume) resting state run was acquired. In the original work, these data were referred to as the “full-k” data in the “scan length” study (Barry et al., 2018). The other functional data also collected in that study were not used here. Physiological data (respiration, pulse) were collected during the session. Pulse data were not used for one subject due to poor quality; both respiration and pulse data were not available for one other subject. These subjects were excluded from fMRI modeling (see section 2.2.4) and following analyses (final included sample: n=, 26.0 ± 4.2 years, 5 F).

### 2.2 Data analysis

The FMRIB Software Library (FSL; version 6.0.3), Spinal Cord Toolbox (SCT; version 6.1), Neptune Toolbox (version 1.211227) (Deshpande & Barry, 2022), AFNI (version 22.0.05), MATLAB (MathWorks, Natick, MA, R2019b), and R (version 4.3.1) were used to perform data analyses (unless otherwise noted).

#### 2.2.1 Motion correction

A mask was manually drawn around the spinal cord and CSF region, which is used to compute a Gaussian weighting mask for motion correction. Note, when using the term “CSF” throughout this paper, this refers to the CSF in the spinal subarachnoid space, unless otherwise stated. Slice-wise motion correction was performed with the Neptune Toolbox and AFNI as previously described (Hemmerling et al., 2023a). Note, one difference is that temporal median filtering was de-selected for the datasets using the Siemens ZOOMit acquisition but retained for the resting state B dataset because it was used as such in the original publication (Barry et al., 2016, 2018).

#### 2.2.2 Noise ROI definition and SpinalCompCor regressor generation

Regressors were derived from noise ROI voxels external to the spinal cord and CSF region. These voxels are expected to contain signal fluctuations from noise sources that similarly affect the timeseries within the cord. Note that noise associated with CSF signal fluctuations was addressed by generating a separate, CSF-specific regressor (see section 2.2.4), as this is a common practice in the literature **(Table S1)**. To create the noise ROI, the mask drawn for motion correction around the spinal cord and CSF was dilated by 18 mm, rounded to the nearest whole voxel (*sct_maths*). This value was chosen based on the dilation to the approximate image boundary for the selective excitation sequences and kept consistent for all datasets. The spinal cord/CSF mask was subtracted from the dilated region, and any voxels at the edge of the slice field-of-view (up to 3 voxels) were removed from the final noise ROI mask (*fslmaths*). Examples of this noise ROI are shown in **Fig. S2**. All voxel timeseries from within the noise ROI were output (*fslmeants*) and arranged in a matrix (rows: time points, columns: voxels). Slice-wise PCA by singular value decomposition was performed on this matrix with MATLAB (*pca*). These steps are depicted schematically **(Fig. 1A-C)**. Components were then selected from the ordered principal components to be SpinalCompCor regressors.

**Fig 1.**
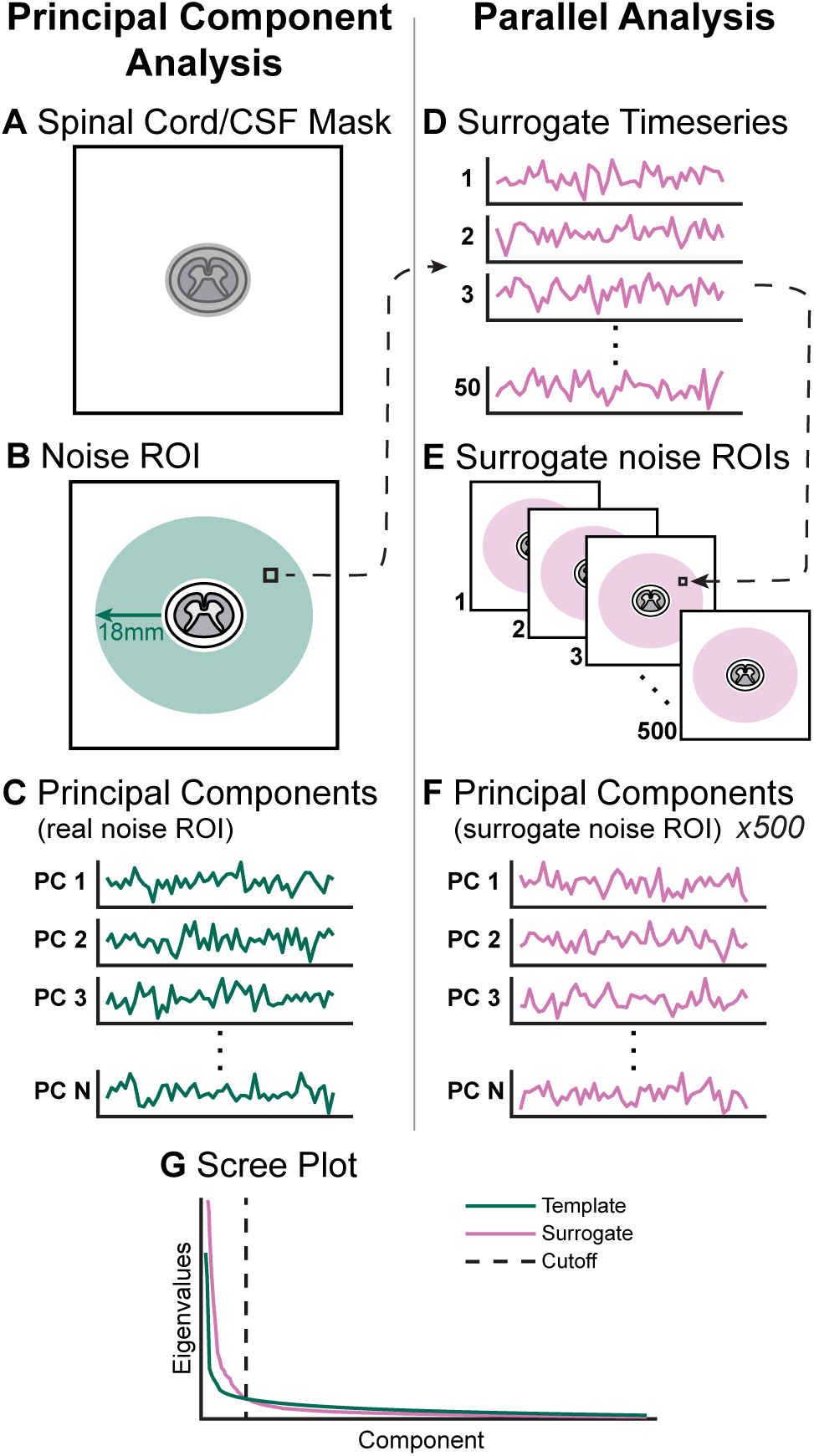
Schematic overview of (left) principal component analysis and (right) parallel analysis methods for each slice. **(A)** The manually drawn spinal cord/CSF contour. **(B)** The noise ROI. **(C)** Slice-wise principal component analysis is performed in the noise ROI, generating N components (N=# Timepoints-1) per imaging slice. **(D)** Surrogate timeseries (50 per voxel) generated from the real voxel timeseries. **(E)** Surrogate slice noise ROI timeseries matrices (500) generated by randomly sampling the surrogate voxel timeseries. **(F)** Principal component analysis is performed in the surrogate slice noise ROI, generating N components. This is repeated for each of the 500 surrogate noise ROIs. **(G)** Scree plot with mean surrogate and template curves; the indicated cutoff is at the intersection of these two curves. Note, the plots shown in this schematic for illustrative purposes and are not actual timeseries or data used in this work.

#### 2.2.3 Surrogate data generation and parallel analysis

To systematically determine how many principal components may be reasonable to retain, a parallel analysis was performed comparing real principal component eigenvalues to those generated from simulated data. An Iterative Amplitude Adjusted Fourier Transform (IAAFT) method was used to generate surrogate timeseries data^1^. Surrogate timeseries share some statistical properties with the original timeseries; in this case, the IAAFT algorithm preserves amplitude and power spectrum traits of the original, but with random phases (Venema et al., 2006). Since the phases are randomly shifted, these surrogate timeseries are, on average, uncorrelated with the original timeseries on which they are based. As such, surrogate timeseries can be used to generate “null” principal components that properly account for temporal dynamics.

For a given template (real) voxel timeseries, an initial surrogate timeseries was first created by shuffling datapoints in the template voxel timeseries. The IAAFT algorithm then iteratively performs spectral adjustment and amplitude adjustment until convergence, or the maximum number of iterations (500) was reached (Venema et al., 2006). This was repeated for each template voxel timeseries in each slice of the noise ROI such that 50 surrogate timeseries were generated from each template (i.e., real) timeseries.

500 surrogate noise ROI timeseries matrices were generated by randomly sampling from the 50 surrogate timeseries for each voxel. Slice-wise PCA was run on the template and each of the 500 surrogate noise ROIs. A scree plot was generated as follows: (1) the template Eigenvalues and mean surrogate Eigenvalues were plotted against the component number; (2) the component number at which these curves intersected was selected as the cutoff number of principal components for that slice. This cutoff represents the last template component that explains greater variance than the surrogate components do by chance. These steps are depicted schematically **(Fig. 1D-G)**. Note, MATLAB version r2022b was used for the parallel analysis because of software availability on the high-performance computing cluster. Northwestern’s high-performance computing cluster, Quest, is equipped with 1231 compute nodes with 80,398 cores and 12.0 petabytes of ESS storage. Nodes used the Red Hat Enterprise Linux 8.12 operating system.

The parallel analysis could be applied to any given dataset to determine a cutoff number of components. The code necessary to generate SpinalCompCor regressors and perform the parallel analysis with IAAFT is provided alongside this publication. Alternatively, the Neptune toolbox for spinal cord fMRI could also be used to perform PCA-based denoising (Deshpande & Barry, 2022).

We implemented the parallel analysis in task and rest fMRI datasets to evaluate whether a different number of components would be designated to retain as SpinalCompCor regressors across these different fMRI acquisitions. Timeseries were first truncated to 160 timepoints to match the resting state A data (the dataset with the fewest timepoints) so that the number of timepoints would not influence the number of principal components in the following parallel analysis. The parallel analysis was then implemented as described above. For each fMRI run, the median cutoff number of components across slices was selected.

#### 2.2.4 Subject-level models with SpinalCompCor regressors

The downstream effect of using SpinalCompCor regressors in subject-level fMRI modeling was tested across all the datasets. The following additional preprocessing was performed (these steps are more thoroughly described in previous work from our lab (Hemmerling et al., 2023a)). First, additional task and nuisance regressors were generated. Physiological data were preprocessed via a bespoke MATLAB script to detect peaks in the respiratory belt, pulse, and CO_2_ traces. An end-tidal CO_2_ (P_ET_CO_2_) breath-hold task regressor (if applicable) and slice-wise RETROICOR (Brooks et al., 2008; Glover et al., 2000) nuisance regressors (8 respiratory and 8 cardiac) were generated from the preprocessed physiological data. A slice-wise CSF regressor was generated from CSF voxels with the top 20% temporal variance (Brooks et al., 2008; Kong et al., 2012). Left and right “%MVC” hand-grasp task regressors were also generated by normalizing the force trace to the participant’s MVC (if applicable). Then, a manual spinal cord mask was drawn on the motion-corrected mean functional image. The high-resolution anatomical image was segmented and registered to the PAM50 template, using labels of the C3 and C7 vertebrae (task and resting state A) or C3 and C5 vertebrae (resting state B) (De Leener et al., 2018; Gros et al., 2019). Then the functional data were also registered with the PAM50 template using the spinal cord mask and initial warp from the anatomical registration.

The subject-level analysis was performed with FSL’s FILM general linear model using the FEAT tool (Jenkinson et al., 2012) (FILM prewhitening; high-pass filter cutoff = 100 s). The models included a subset of task (P_ET_CO_2_ or Left/Right %MVC for the breath-hold and hand-grasp task, respectively) and nuisance regressors (X and Y motion (MOCO), CSF, 8 cardiac and 8 respiratory RETROICOR (RETRO), and SpinalCompCor (PCA)). A Right Grasp > 0 and Left Grasp > 0 contrast were defined for the hand-grasp task; a P_ET_CO_2_ > 0 contrast was defined for the breath-hold task. The resting state models were computed identically except without task regressors.

Three models were computed for each dataset:

1. “Base” model with no SpinalCompCor components (19 nuisance regressors)

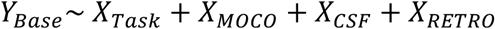
2. “Extended” model with all Base model and additional PCA-derived regressors (>19 nuisance regressors)

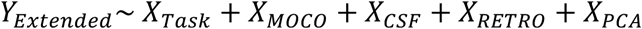
3. “SpinalCompCor” model *without* physiological recording-derived regressors (i.e., RETROICOR) and with PCA-derived regressors (>3 nuisance regressors)

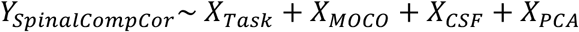

The median number of principal components chosen by the parallel analysis across slices was selected for each dataset to avoid creating models with different numbers of regressors in each slice. Subject-level models were computed on the full timeseries, so the parallel analysis was run again on the full timeseries before selecting the number of component regressors for modeling. Models were only computed for voxels within a spinal cord mask to minimize computational burden. For a few representative fMRI runs, the GLM was rerun without a mask (i.e., in the entire field of view) so that spatial information about the principal component regressors could be examined outside of the cord.

#### 2.2.5. Evaluation of nested models

Next, we addressed whether adding principal component noise regressors derived from outside the spinal cord successfully explained variance inside the spinal cord. The second model (Extended) was selected to examine whether the PCA-derived regressors are beneficial *in addition* to standard denoising. Thus, a voxel-wise *F*-test was performed between the full (Extended) and reduced (Base) models. PAM50 template gray matter and white matter masks were warped to the native space for each run, thresholded (at 0.5), and binarized to provide subject-specific tissue masks. Median *F*-statistics and the proportion of statistically significant voxels, as determined by the *F*-test (*p*<0.05, not corrected for multiple comparisons), were computed in these masks.

Considering the potential spinal cord segment-dependent effect observed in the *F*-test results, we wanted to evaluate whether there was in fact a segment-dependent benefit to using the Extended model compared to the Base model. We used a mixed effects generalized additive model with a quasibinomial family and logit link function, which was fit using restricted maximum likelihood (REML). The dependent variable, proportion of significant voxels, was regressed on spinal cord level, task, and tissue type as fixed effects, and subject was included as a random effect, saving the parameter estimate associated with spinal cord level as the test statistic. Spinal cord level was coded as −1, 0, and +1 for C6, C7, and C8, respectively, for hand-grasp, breath-hold, and resting state A datasets, or as –1, 0, and +1 for C4, C5, and C6, respectively, for the resting state B dataset. To build a null distribution, we relied on permutation testing (5000 permutations). Within each permutation, each subject’s spinal cord levels were randomly “flipped” (rostrocaudal vs. caudorostral) via sign flipping (*e.g*., <u>−1, 0, and +1 *or* +1, 0,</u> <u>and −1 for C6, C7, and C8, respectively)</u>, which would preserve within-participant autocorrelation across adjacent levels. The spinal cord level parameter estimate (test statistic) from each permutation was saved, and the observed test statistic was compared to this null distribution to calculate a *P*-value.

#### 2.2.6. Evaluation of recording-based vs. data-driven regressors

The SpinalCompCor model represents a scenario in which high-quality physiological traces may not be available to generate nuisance regressors, such as RETROICOR regressors. Here, it is worthwhile to investigate whether SpinalCompCor regressors could be used as a sufficient alternative to recording-based RETROICOR regressors. In addition to the task regressor contrasts stated above, *F*-tests were set up in FSL by first defining a contrast for each RETROICOR regressor (Base model) or each SpinalCompCor regressor (SpinalCompCor model) and then entering all of those contrasts into an omnibus *F*-test for each respective model.

We then determined whether the set of RETROICOR regressors in the Base model explained variance in a spatially similar way to the SpinalCompCor regressors in the SpinalCompCor model. A spatial correlation (Spearman) between the output *z*-score transformed *F*-statistics for each of those *F*-tests was computed (in native subject space).

#### 2.2.7. Temporal signal-to-noise ratio (tSNR)

To evaluate the overall effectiveness of these denoising procedures, the tSNR was calculated for each model. The functional mean was added back to the model residuals, and then the tSNR was calculated (mean / standard deviation). Within each dataset, the mean tSNR was compared between models (Base, Extended, and SpinalCompCor) using pairwise *t*-tests (two-sided, paired, with Bonferroni correction) using the *rstatix* package in R.

#### 2.2.8. Group modeling and comparisons

The hand-grasp and breath-hold task datasets were carried through to group-level activation mapping. For this analysis, each participant’s second fMRI run was also considered (i.e., the parallel analysis and subject-level modeling described above were also performed on these runs). To assess repeatability across the two runs, an interclass correlation coefficient (ICC) was calculated in R using a two-way random effects model, single rating, and absolute agreement with the icc() function from the irr package. Parameter estimate maps associated with each run were averaged for each subject and then concatenated across subjects. For group-level analysis, a non-parametric one-sample *t*-test with threshold-free cluster enhancement was performed with 5000 permutations (*randomise*) (Nichols & Holmes, 2002; Smith & Nichols, 2009; Winkler et al., 2014). This was performed for each model (Base, Extended, SpinalCompCor) and contrast. The proportion of voxels that were “Now active” and “No longer active” with the Extended model compared to the Base model and with the SpinalCompCor model compared to the Base model were extracted in ROI masks. A Dice similarity coefficient (DSC) was used to assess the similarity of the spatial distribution of significant voxels between models (*3ddot*).

#### 2.2.9. Functional connectivity

The denoised resting state fMRI datasets (i.e., the residuals from the subject-level modeling) were carried through to a seed-based functional connectivity analysis. ROIs were defined as the left-ventral, left-dorsal, right-ventral, and right-dorsal horns from the PAM50 atlas. The ROI masks were warped to each subject’s native space, thresholded (at 0.5), and binarized.

Slicewise correlation matrices were calculated using the mean timeseries in each ROI and were Fisher r-to-z-transformed (*3dNetCorr*). A full correlation was used (Barry et al., 2018). The mean of the ventral-ventral (V-V), dorsal-dorsal (D-D), ventral-dorsal within hemicord (V-D Within), and ventral-dorsal between hemicords (V-D Between) correlations were averaged across slices (Kaptan et al., 2023; Weber et al., 2018). For each resting state dataset and correlation type (V-V, D-D, V-D Within, V-D Between), the mean correlation was compared between models (Base, Extended, and SpinalCompCor) using pairwise *t*-tests (two-sided, paired, with Bonferroni correction) using the *rstatix* package in R.

## 3 RESULTS

### 3.1 Parallel analysis

A parallel analysis was run to determine the number of principal components that should be considered as regressors for fMRI modeling. This analysis generated a scree plot for each slice of every fMRI run, from which the cutoff number of principal components was recorded (see **Fig. 1G** as an example). The median cutoff number of principal components across slices is shown in **Fig. 2A** for all four datasets, truncated to match the number of volumes. The distribution across slices shows that the cutoff number of components is relatively consistent across most subject/slices **(Fig. 2B)**. However, there are areas of higher variability, e.g., in the inferior-most slices for the hand-grasp task and resting state A datasets. It is pertinent to note that three datasets (breath-hold, hand-grasp, resting state A) were acquired on a Siemens scanner using a sequence with selective excitation in an inner FOV with a TR of 2s. Selective excitation should lessen susceptibility artifacts in areas near air cavities (Rieseberg et al., 2002), such as the spinal cord. The resting state B fMRI data were acquired on a Philips scanner, using a 3D acquisition, did not use a sequence with selective excitation, and had a VAT of 2.76s. While all data were acquired in the cervical cord, the resting state B imaging volume was also centered more rostrally (approximately vertebral levels C2-C5) than the other acquisitions (approximately vertebral levels C4-C7), which may impact the specific noise characteristics captured by PCA.

**Fig. 2.**
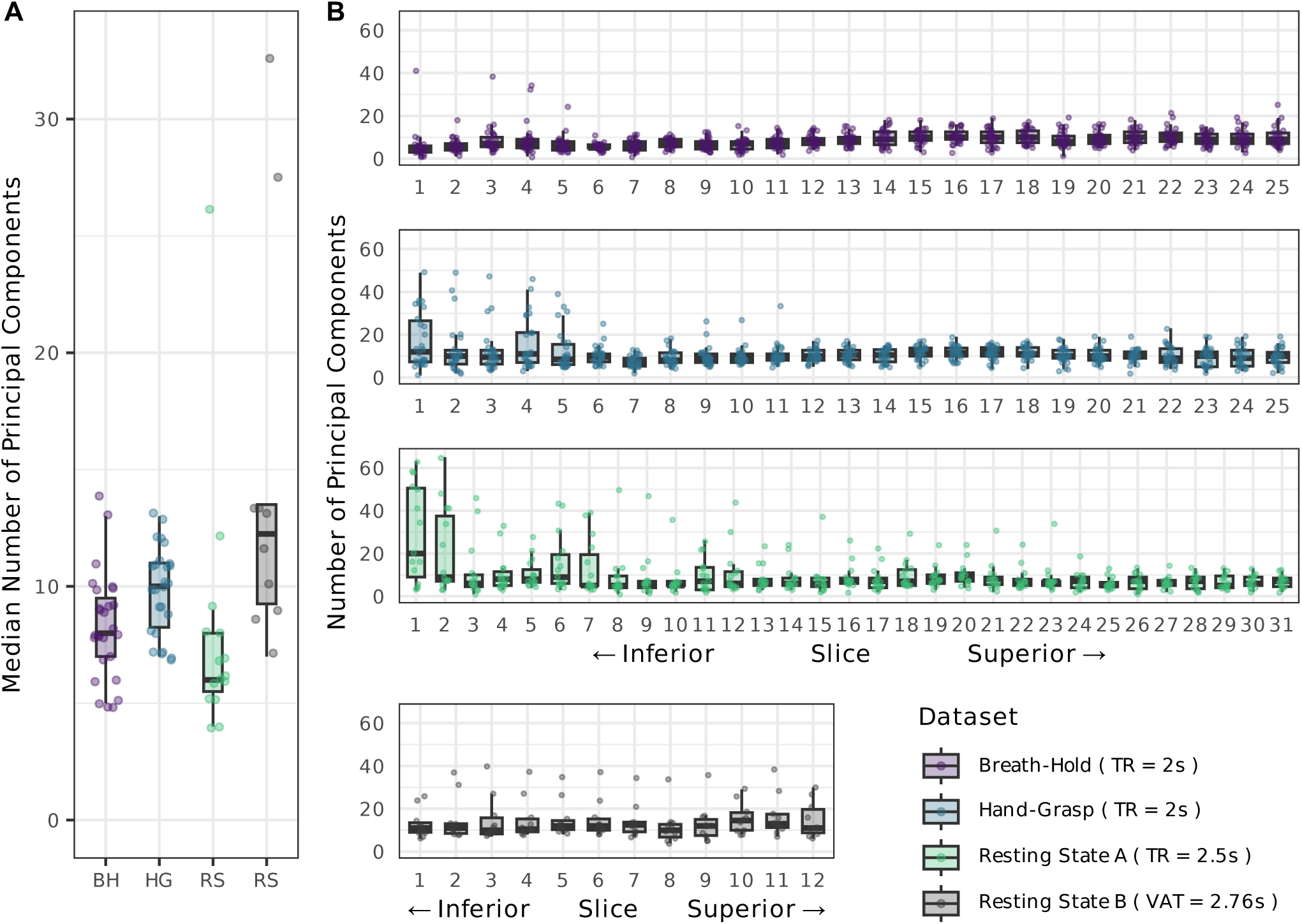
Parallel analysis summary across datasets. **(A)** Median cutoff number of principal components for each fMRI run (each datapoint represents the parallel analysis result across slices for one subject’s fMRI run). **(B)** The cutoff number of principal components in each subject and slice. Note, the parallel analysis was performed on data restricted to the same number of timepoints (i.e., fMRI data that were originally longer than 160 timepoints were truncated).

Longer fMRI runs were truncated to 160 timepoints for this parallel analysis, so the PCA performed here was not biased by the scan length. Having more timepoints does, however, influence the cutoff number of principal components selected in the parallel analysis. For example, the hand-grasp task dataset’s original length is 300 timepoints, and truncating the dataset to 160 timepoints decreases the median number of cutoff components for nearly all subjects’ scans **(Fig. S3)**.

There are also two notable outliers in the median cutoff number of principal components for the resting state B dataset. To explore this, the median number of principal components was also compared to the in-plane framewise displacement **(Fig. S4A)**. We see a significant negative trend between number of components and overall motion across all datasets (*r*=-0.75, *p*=0.012) and observe that the two outlier scans with the highest median number of principal components also had some of the lowest overall motion. Motion may also be positively associated with the magnitude of the eigenvalue associated with the first principal component, and **Fig. S4B** shows that these outliers have the lowest first eigenvalue. However, the positive correlation between these elements is non-significant (*r*=0.25, *p*=0.49).

### 3.2 Spatial information from component regressors

Maps of the fitted parameter estimates show how variance associated with each SpinalCompCor regressor was fit across the entire cord and neck region. For the datasets in which the subject-level model was rerun on the entire field of view, stronger parameter estimates were observed in a few common areas: neck muscle, blood vessels, and CSF. Examples from hand-grasp scans are shown in **Fig. 3** (A: blood vessel/CSF; B: blood vessel/CSF; C: no predominant area; D: neck muscle). Parameter estimates from panels A and B overlaid on mean images are provided in **Fig. S5**, showing the location of stronger parameter estimates with respect to the CSF and white matter. Examples are also provided from a breath-hold scan in **Fig. S6**, from a resting state A scan in **Fig. S7**, and from a resting state B scan in **Fig. S8**. All parameter estimate maps shown are from the Extended model, but parameter estimate maps for the same principal component regressors in the SpinalCompCor model are nearly identical.

**Fig. 3.**
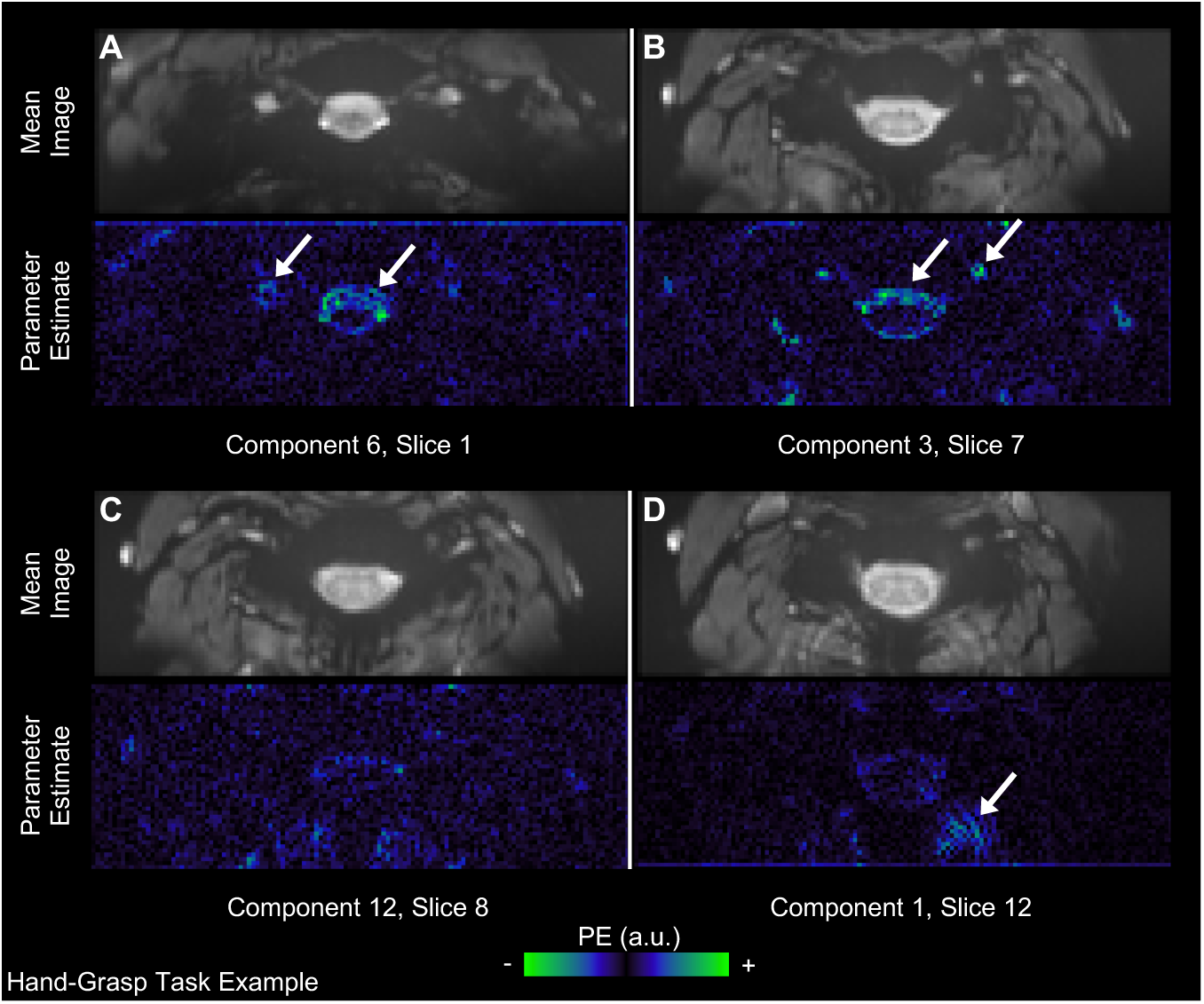
Parameter estimate fits for SpinalCompCor regressors in an example subject from the hand-grasp task dataset. The motion corrected mean image and corresponding principal component parameter estimate maps are shown for each example component/slice. Arrows point out stronger parameter estimates in areas with neck muscle, blood vessels, and CSF. There are 25 slices in this acquisition, numbered 0-24. (PE=parameter estimate, a.u.=arbitrary units)

### 3.3 Evaluation of nested models

The Base and Extended models were calculated for each breath-hold, hand-grasp, and resting state fMRI run to explore the effect of adding SpinalCompCor regressors in addition to base denoising. Differences between nested Base and Extended models (i.e., the reduced and full models, respectively) were evaluated with a voxel-wise nested *F*-test. Summary metrics were extracted from gray matter and white matter ROIs in spinal cord segments in the native subject space (within tissue masks warped from the PAM50 template). Specifically, the median *F*-statistic and the proportion of statistically significant voxels (*F*-test) were calculated in ROIs for each fMRI run **(Fig. 4)**.

**Fig. 4.**
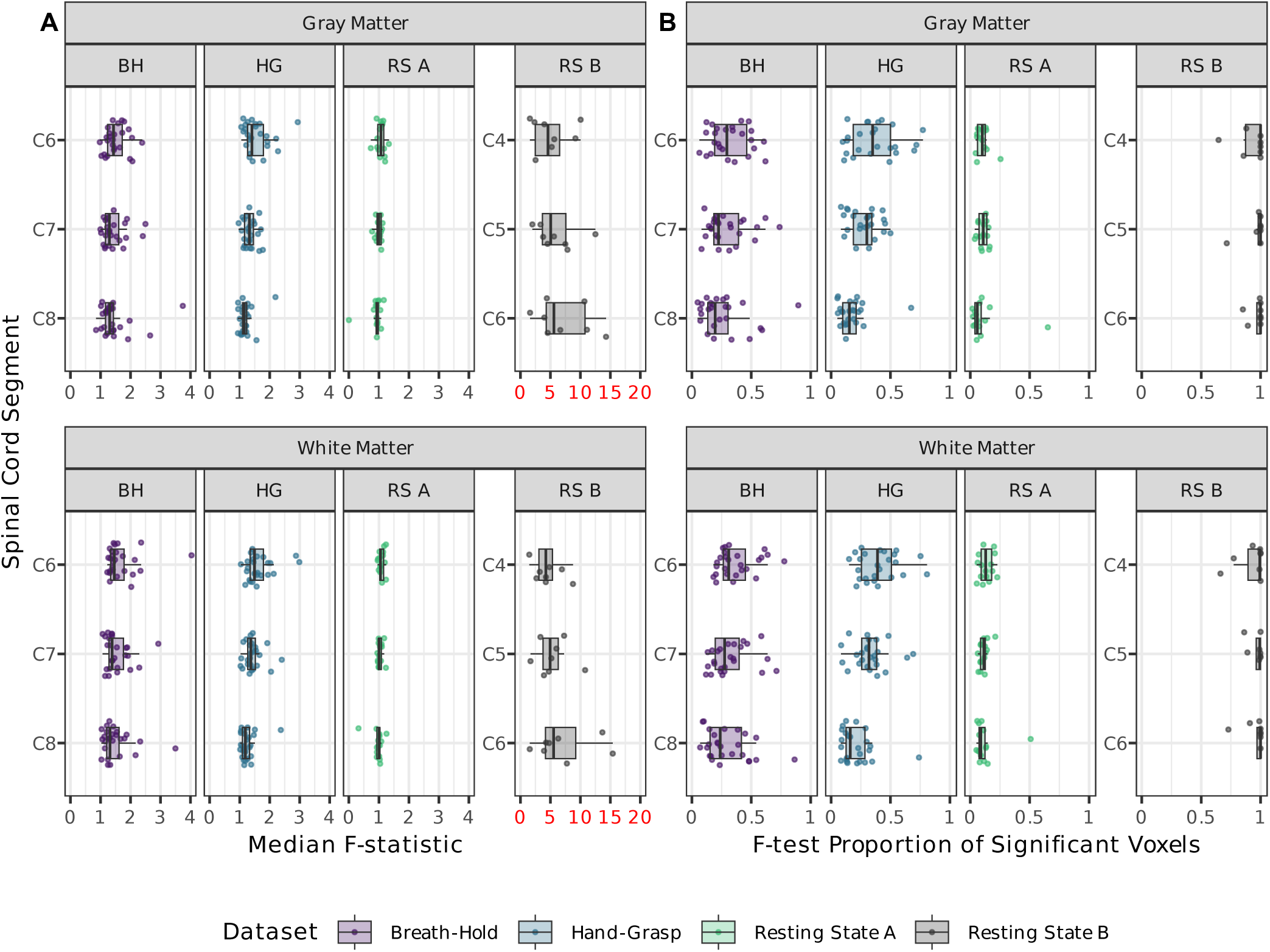
Summary of F-test comparing the nested Base and Extended models. Summary statistics are shown in ROIs of gray matter and white matter across spinal cord segments C6-C8 or C4-C6. **(A)** Median F-statistic in each of the spinal cord ROIs. **(B)** The proportion of voxels in the ROI for which the Extended model significantly fit additional variance compared to the Base model (p<0.05, not corrected for multiple comparisons). Each datapoint represents one fMRI run. Resting state B median F-statistic axis is colored red to draw attention to the scaling.

For both the breath-hold and hand-grasp task datasets, the median *F*-statistic appears to decrease slightly from superior to inferior (C6 to C8) in both gray and white matter. The proportion of statistically significant voxels in the *F*-test similarly appears to decrease from superior to inferior in both gray and white matter. In the resting state B dataset, the percent of significant voxels (*F*-test) clusters near 100% and may also differ across levels. Assessing this using a mixed effects model and flipping the spinal cord segments, we can reject the null that all spinal cord segments benefit identically (p < 0.0002).

### 3.4 Evaluation of recording-based vs. data-driven regressors

In the absence of high-quality physiological traces to derive nuisance regressors, such as RETROICOR regressors, data-driven regressors may be a desirable substitute. *F*-tests for the set of RETROICOR regressors in the Base model compared to the set of SpinalCompCor regressors in the SpinalCompCor model are compared using a spatial correlation between *F*-statistic maps **(Fig. 5)**. This spatial correlation illustrates how consistently each set of nuisance regressors accounts for variance in the BOLD signal across the spinal cord. The breath-hold task, hand-grasp task, and resting state A datasets have a moderate correlation, while the resting state B dataset has a low correlation. Thus, it is likely that both the RETROICOR and SpinalCompCor regressors capture physiological noise, although these models do not capture them equivalently.

**Fig. 5.**
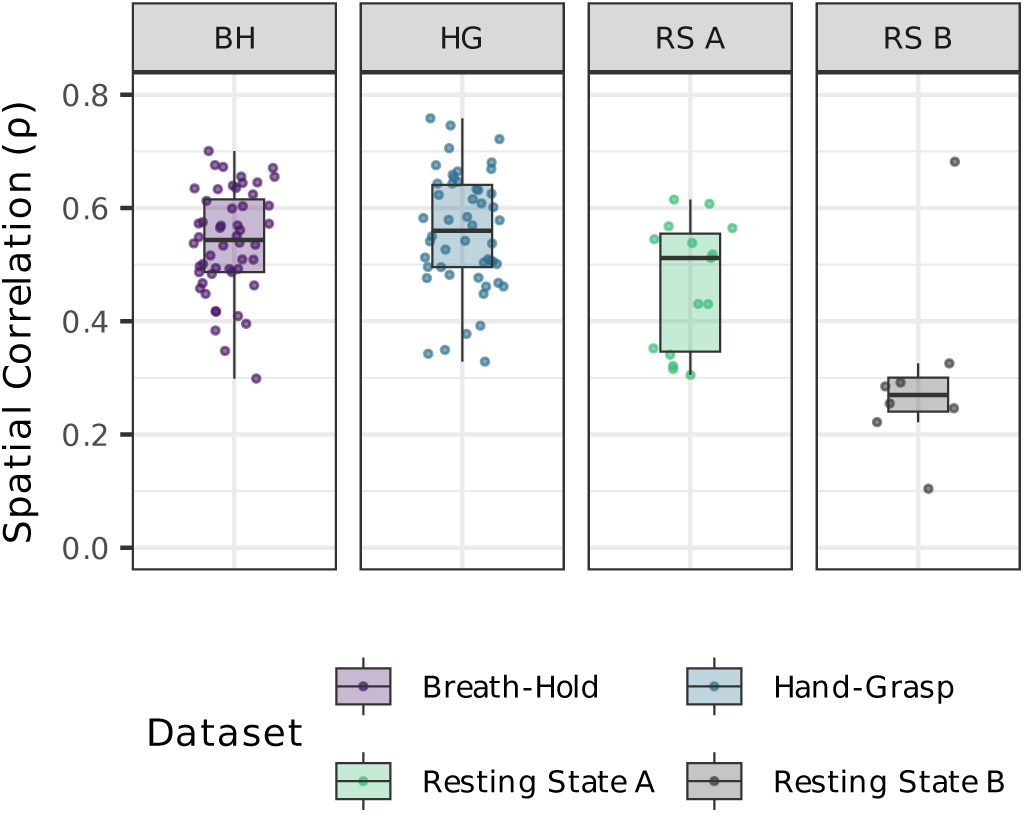
Spatial similarity of recording-based versus data-driven nuisance regressor F-statistic maps in each dataset. Voxel-wise Spearman correlation coefficients between F-statistic maps for each run (in spinal cord voxels). The F-statistic maps result from omnibus F-tests performed separately on the set of RETROICOR regressors (Base model) and the set of SpinalCompCor regressors (SpinalCompCor model).

### 3.5 tSNR

The tSNR after denoising was calculated for each dataset and model **(Fig. 6)**. The Extended model, which contains both the RETROICOR and SpinalCompCor regressors and should, by definition, explain the greatest variance, significantly improves the tSNR of the residual timeseries in all four datasets, as compared to the Base or SpinalCompCor only models. The tSNR of the Base and SpinalCompCor models shows a statistically significant difference in three out of four datasets, indicating that SpinalCompCor regressors may not be a perfect substitute for physiological recording-based regressors. Combined with the results in Figure 5, this suggests that SpinalCompCor may only partially mitigate the problem of physiological noise in datasets without available physiological recordings.

**Fig. 6.**
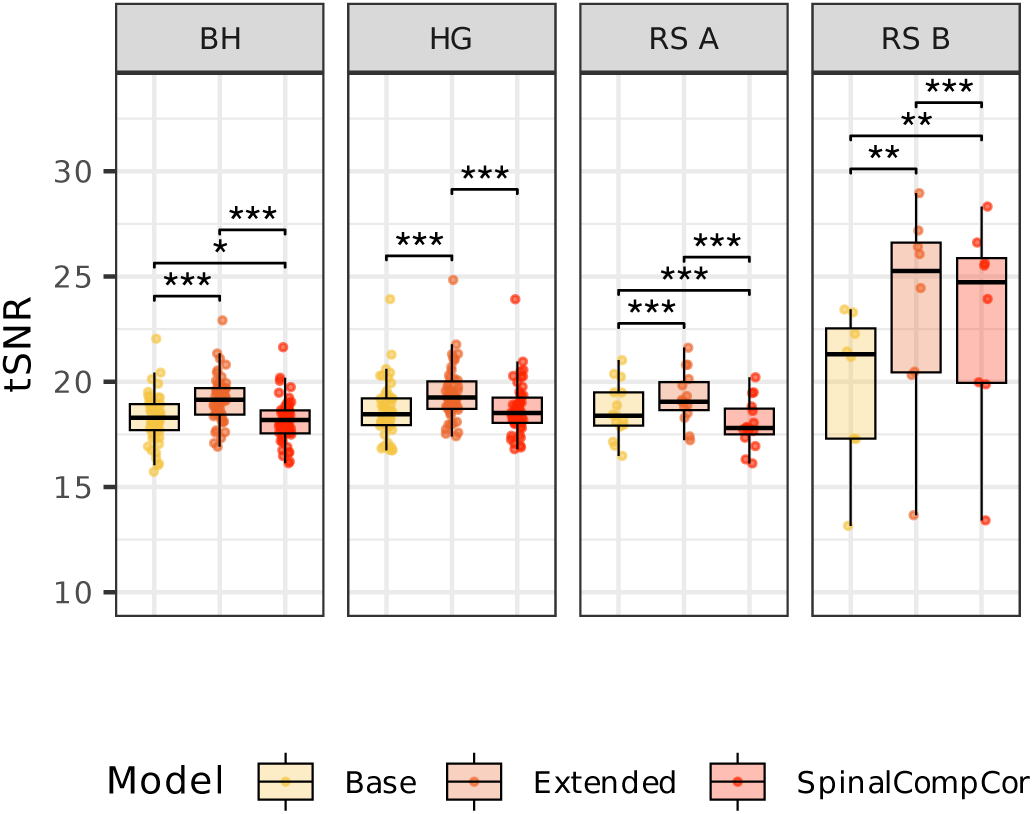
Spinal cord tSNR across Base, Extended, and SpinalCompCor (only) models for all four datasets. (t-tests, p < 0.05*, p < 0.01**, p < 0.001***; Bonferroni corrected).

**Table 1.** Spinal cord tSNR between Base, Extended, and SpinalCompCor (only) models.

|  | Model Comparison | Mean Difference [95% CI] | t | p |
| --- | --- | --- | --- | --- |
| **BH** | Base vs. Extended | -0.895 [-0.964, -0.826] | -26.048 | **< .001** |
|  | Base vs. SpinalCompCor | 0.102 [0.030, 0.173] | 2.852 | **0.019** |
|  | Extended vs. SpinalCompCor | 0.997 [0.971, 1.023] | 76.170 | **< .001** |

|  | Model Comparison | Mean Difference [95% CI] | *t* | *p* |
| --- | --- | --- | --- | --- |
| **HG** | Base vs. Extended | -0.775 [-0.834, -0.716] | -26.483 | **< .001** |
|  | Base vs. SpinalCompCor | -0.064 [-0.124, -0.003] | -2.117 | 0.118 |
|  | Extended vs. SpinalCompCor | 0.711 [0.690, 0.733] | 65.706 | **< .001** |
| **RS A** | Base vs. Extended | -0.654 [-0.876, -0.431] | -6.295 | **< .001** |
|  | Base vs. SpinalCompCor | 0.591 [0.370, 0.812] | 5.731 | **< .001** |
|  | Extended vs. SpinalCompCor | 1.244 [1.188, 1.301] | 47.241 | **< .001** |
| **RS B** | Base vs. Extended | -3.520 [-5.112, -1.928] | -5.227 | **0.004** |
|  | Base vs. SpinalCompCor | -2.988 [-4.507, -1.469] | -4.651 | **0.007** |
|  | Extended vs. SpinalCompCor | 0.532 [0.402, 0.661] | 9.688 | **< .001** |

Pairwise *t*-test results for the mean tSNR between each model across all four datasets. The *p*-values were adjusted for multiple comparisons using Bonferroni correction. CI is the confidence interval, *t* is the *t*-statistic, *p* is the adjusted *p*-value (bold font if *p* < 0.05).

### 3.6 Group analysis

A group analysis was performed on the hand-grasp task and breath-hold task data to test the effect of SpinalCompCor regressors on group-level activation, comparing results between the Base and Extended models and between the Base and SpinalCompCor models. Example axial slices of the hand-grasp activation map are shown for the Left>0 and Right>0 contrasts for the Base, Extended, and SpinalCompCor models **(Fig. 7A,B,D)**. To directly compare the spatial distribution of the difference in activation between the Base and the other models, the voxels that changed between conditions were mapped and summarized across the level and tissue ROIs **(Fig. 7C,E and Fig. S9)**. These panels show voxels that were not active for the Base model but are now active for the Extended or SpinalCompCor model (blue), voxels which were active for the Base model but are no longer active for the Extended or SpinalCompCor model (red), and voxels that were active for both (gray). A higher proportion of voxels in the dorsal gray matter are now active with the Extended or SpinalCompCor model, while the opposite is seen for the ventral gray matter. Often, the voxels that change between conditions are at the edge of a cluster of active voxels. Of note, “now active” voxels are observed in the gray matter intermediate zone, suggesting that the addition of SpinalCompCor regressors was necessary to detect this focal effect.

**Fig. 7.**
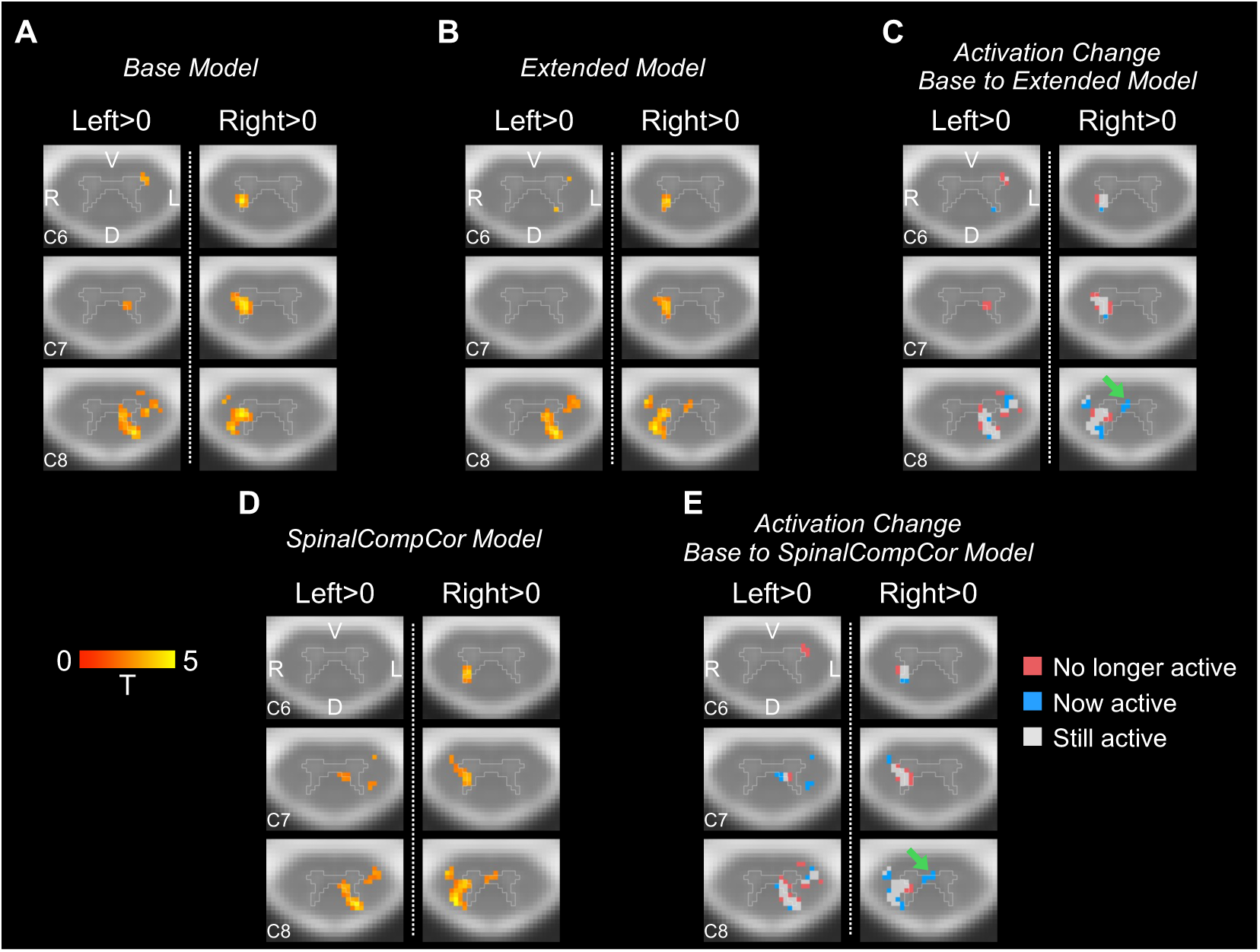
Comparison of Base vs. Extended and Base vs. SpinalCompCor model group analysis for the hand-grasping task. Example activation map axial slices are shown for Left>0 and Right>0, for the **(A)** Base model and **(B)** Extended model. **(C)** Example axial slices of the activation change for the Base to Extended models. Example axial slices are also shown for the **(D)** SpinalCompCor model activation map and for the **(E)** Base to SpinalCompCor activation change. Green arrows point at voxels in the intermediate zone that are now active. Activation maps show significant t-statistics (p<0.05, Family-wise error rate corrected). Axial slices are shown in radiological view/orientation and have an outline of the PAM50 gray matter mask overlaid. The same three slices are shown for panels A-E. (V=ventral, D=dorsal, R=right, L=left).

Similarly, “activation” maps for the breath-holding task group analysis are shown for the Base, Extended, and SpinalCompCor models **(Fig. 8A,B,D)**. The difference in activation is also shown between the Base and Extended or SpinalCompCor models as the activation change **(Fig. 8C,E and Fig. S10)**. Approximately 7-19% of the ventral gray matter voxels are no longer active for the Extended or SpinalCompCor model. Voxels that had a change in condition are also primarily at the edges of a cluster of active voxels.

**Fig. 8.**
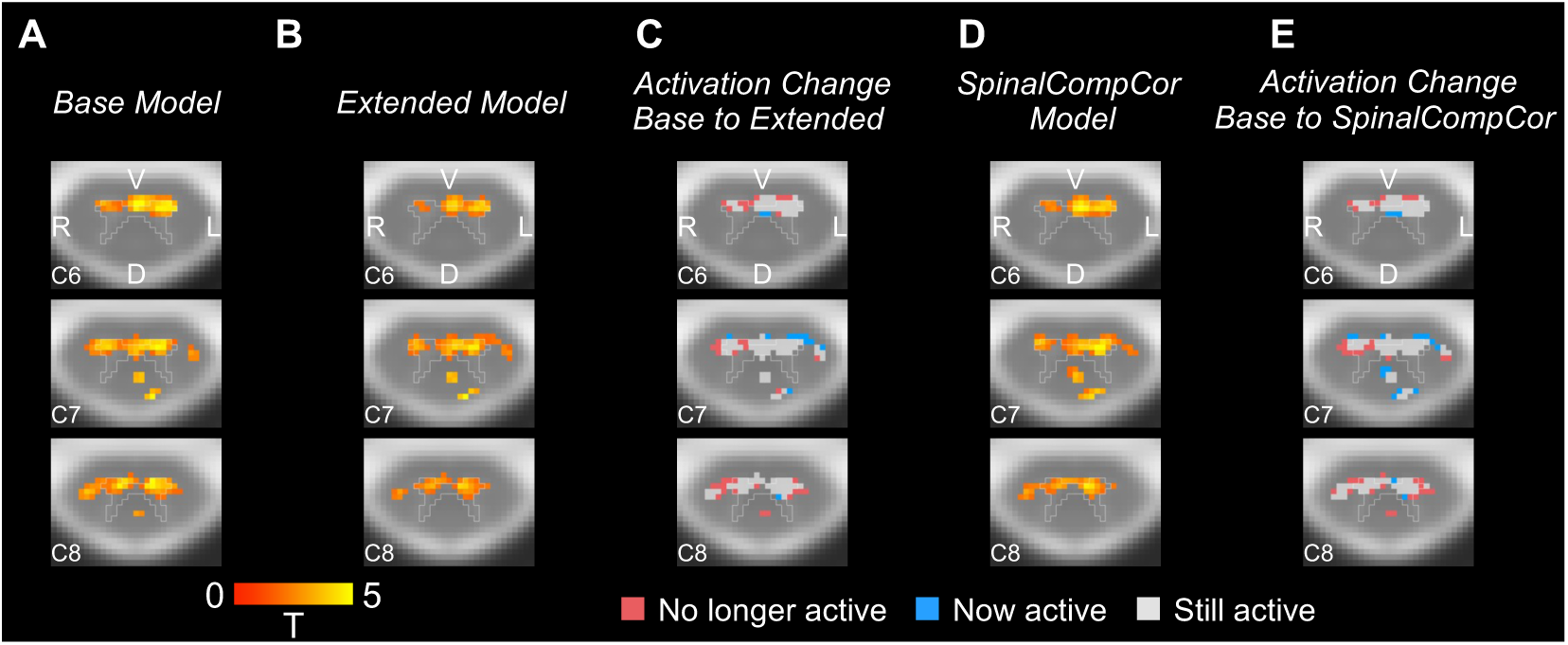
Comparison of Base vs. Extended and Base vs. SpinalCompCor model group analysis for the breath-holding task. Example activation map axial slices are shown for the **(A)** Base model and **(B)** Extended model. **(C)** Example axial slices of the activation change for the Base to Extended models. Example axial slices are also shown for the **(D)** SpinalCompCor model activation map and for the **(E)** Base to SpinalCompCor activation change. Activation maps show significant t-statistics (p<0.05, Family-wise error rate corrected). Axial slices are shown in radiological view/orientation and have an outline of the PAM50 gray matter mask overlaid. The same three slices are shown for panels A-E. (V=ventral, D=dorsal, R=right, L=left).

The total number of significant voxels and distribution of *t*-statistics in significant voxels for each dataset and model is shown **(Fig. S11)**, with more similar distributions between the Extended and SpinalCompCor models across datasets. The spatial similarity of activation between models was also assessed with a DSC **(Fig. S12)**, showing that the area of activation for the Extended and SpinalCompCor model to be the most similar for both tasks, although modestly. This indicates that the sets of SpinalCompCor regressors included in each of these models could be driving this higher similarity.

### 3.7 Functional connectivity

Functional connectivity estimates for both resting state fMRI datasets were computed **(Fig. 9)**. In the resting state B dataset, the Base model has significantly higher correlations than the Extended and SpinalCompCor models for all comparisons, which could represent the removal of spurious noise-driven correlations or true underlying correlations between ROIs with the use of SpinalCompCor regressors in denoising **(Table 2)**. Note that some differences do exist between our processing methods and connectivity quantification from the approach in the prior literature, resulting in some differences in these connectivity estimates and the original papers (Barry et al., 2018; Weber et al., 2018).

**Fig. 9.**
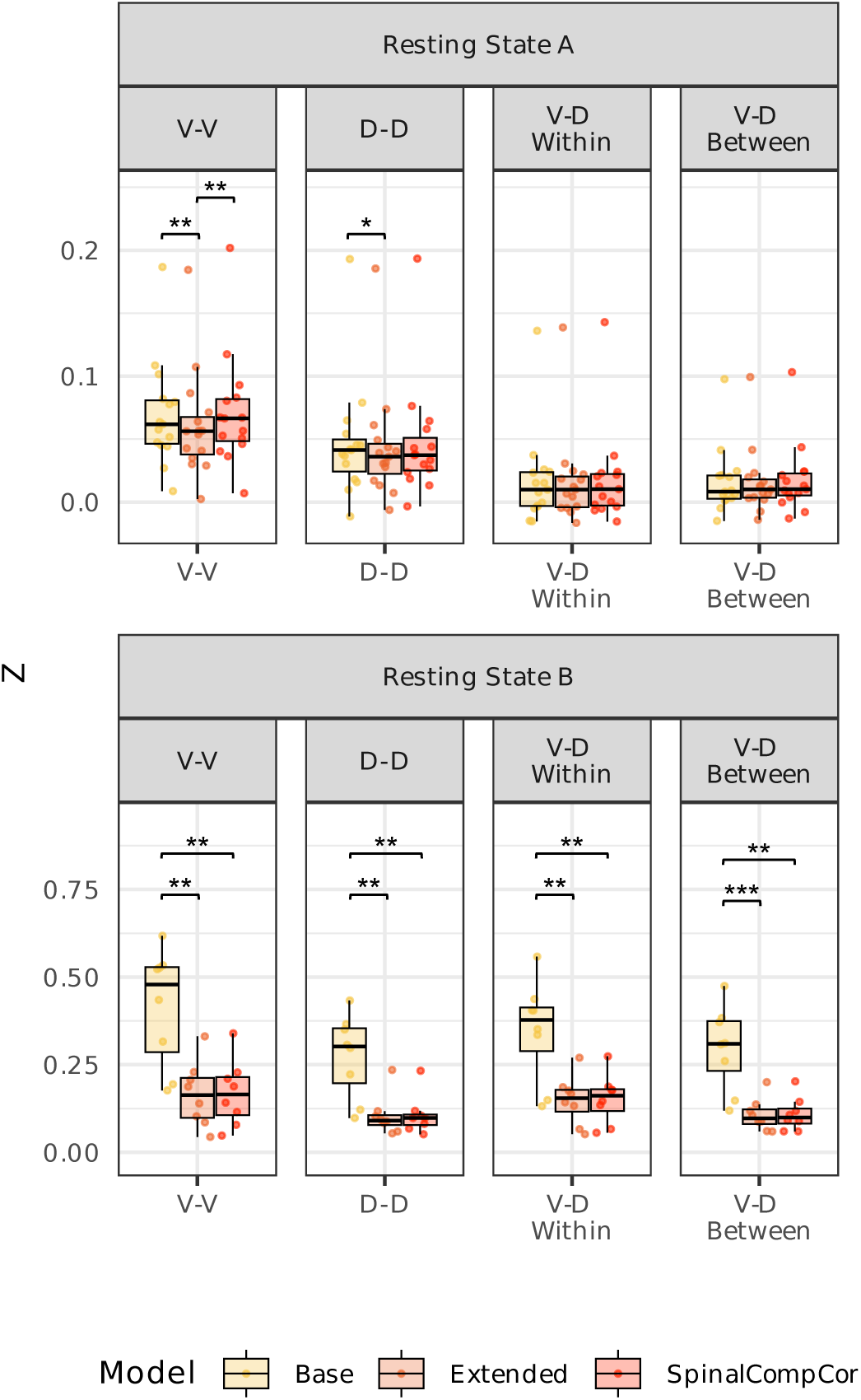
Resting state functional connectivity across subjects for the (top) Resting State A and (bottom) Resting State B datasets for the Base, Extended, and SpinalCompCor models. Average correlation (Fisher r-to-z transformed) between ventral horns (V-V), dorsal horns (D-D), ventral to dorsal horns within the same hemicord (V-D Within), ventral to dorsal horns between hemicords (V-D Between). (t-tests, p < 0.05*, p < 0.01**, p < 0.001***; Bonferroni corrected).

**Table 2.**
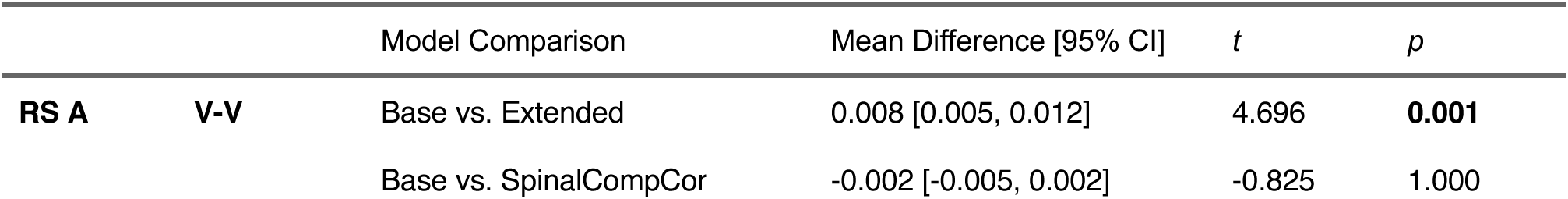
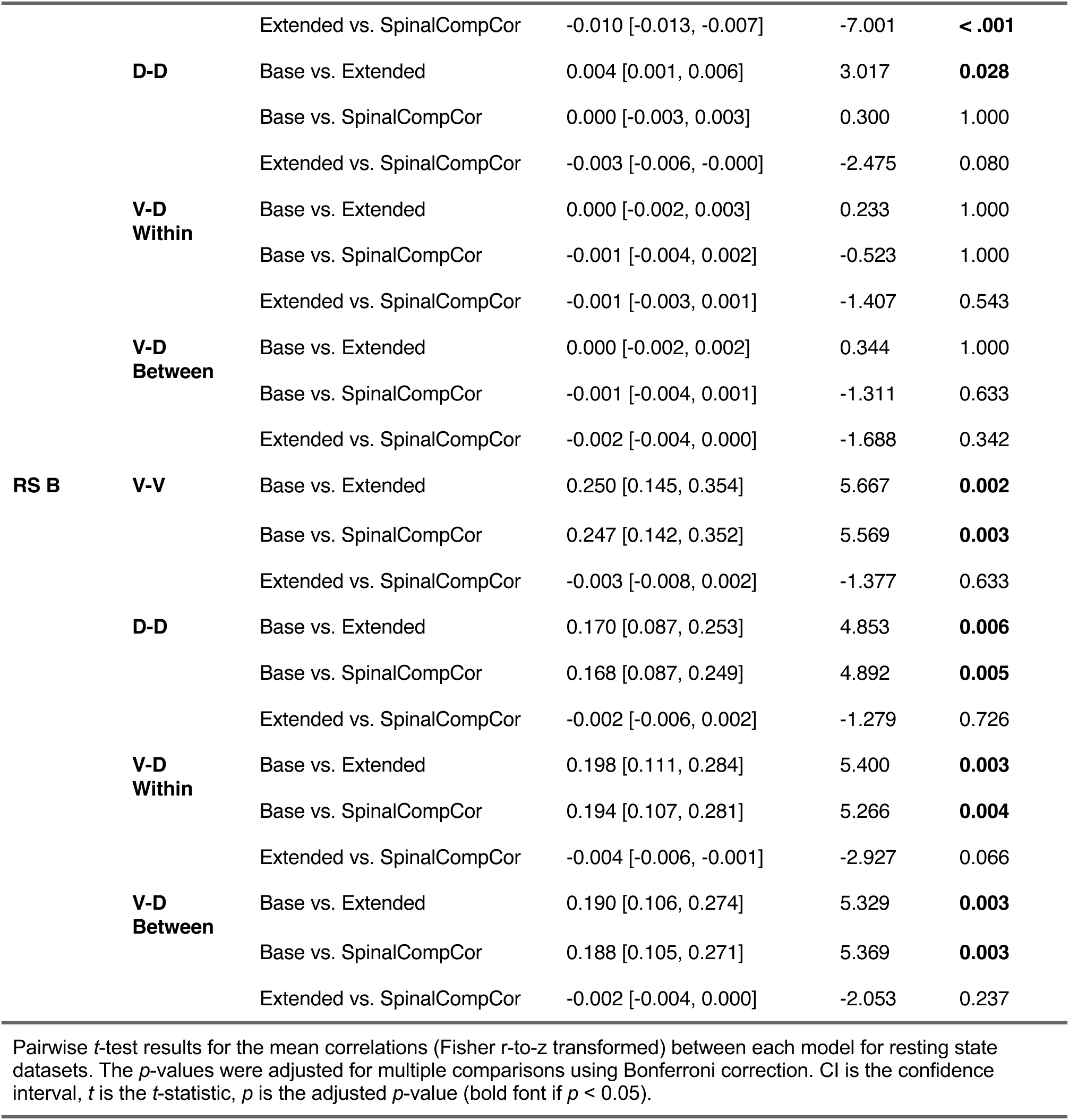
Resting state functional connectivity between Base, Extended, and SpinalCompCor (only) models.

## 4 DISCUSSION

### 4.1 Data-driven denoising in spinal cord fMRI

In this study, we formalize a framework for performing PCA of noise ROI timeseries in spinal cord fMRI data to derive nuisance regressors, using a parallel analysis to automatically determine the appropriate number of component-based nuisance regressors to avoid overfitting. Our work builds on prior research methods developed to denoise both brain and spinal cord fMRI timeseries. In these prior studies, PCA-based nuisance regression in spinal cord fMRI has been implemented but not specifically evaluated, and we lack a standard approach for defining the number of nuisance regressors to be incorporated into the timeseries model. Thus, the current work specifically assesses PCA-based denoising in the spinal cord (i.e., SpinalCompCor), providing a standard, automated approach for deciding the number of PCA-based nuisance regressors to include, and considers the impact of such denoising on the mapping of task activation.

To begin, there are key methodological similarities and differences between the implementation of PCA-based denoising of spinal cord fMRI data in past literature and the SpinalCompCor implementation in the present work that should be discussed. First, Barry et al. (2014) and Barry et al. (2016) define a noise ROI by logically inverting a mask drawn around the entire spinal cord and CSF region. Based on these studies, the Neptune toolbox for spinal cord fMRI analysis contains methods for ‘non-cord’ PCA-based noise regression (Deshpande & Barry, 2022). Similarly, another study derives regressors with PCA from CSF and ‘not-spine’ ROIs and then regresses them from the resting state fMRI timeseries along with other noise regressors (RETROICOR and motion parameters) (Combes et al., 2023). One other resting state fMRI study also includes PCA-derived CSF regressors in their noise model (Landelle et al., 2023). Here, we opt to limit PCA to a noise ROI that does not include the spinal canal CSF, instead addressing pulsatile CSF artifacts with a dedicated regressor. In recent spinal cord fMRI literature, this noise is most often addressed with a dedicated CSF regressor, whether derived by signal averaging or via CSF-only PCA methods **(Table S1)**. Our goal was to determine whether the SpinalCompCor method is beneficial beyond the base denoising that is standard in the field. Whether to include spinal canal CSF in the noise ROI is a nuanced choice that could be addressed in future work.

Furthermore, in the Barry et al. 2016 method, components were selected for each slice based on their associated variance. Specifically, the components retained represented up to 80% of the cumulative variance, or where the difference in variance explained between successive components was less than 5% (Barry et al., 2016), resulting in a different number of components for each slice. In the current work, principal components are retained based on a parallel analysis, which compares template (real data) eigenvalues to surrogate (simulated data) eigenvalues to determine a cutoff number of components for each slice, typically around 9 components in the data considered here (see next section for further details) **(Fig. 2)**. Different thresholds may be needed for different fMRI datasets, and the parallel analysis automatically accounts for this variability by finding the number of components that explain greater variance than the surrogates do by chance. The median across slices is then selected as the number of components to retain as SpinalCompCor regressors. This final step avoids a difference across slices in the number of regressors and model degrees of freedom, simplifying the analysis but potentially over or underestimating the number of regressors that “should” be used for each slice.

Finally, the Barry et al. method regresses the PCA-derived nuisance regressors from the fMRI timeseries prior to motion correction to improve the efficacy of volume registration. In the current work, however, we incorporated the SpinalCompCor regressors into the subject-level GLM, opting instead to minimize the number of regression steps in the analysis pipeline. Although this eliminates the potential benefits to motion correction described in the Barry et al. method, this choice should simplify preprocessing and avoids the possible reintroduction of artifacts back into the data that can occur with sequential regression steps (Lindquist et al., 2019).

### 4.2 Selecting principal components as nuisance regressors

A parallel analysis with surrogate timeseries data was used to systematically determine how many principal components to retain, to minimize overfitting BOLD signal variance. This analysis outputs a cutoff number of principal components for each slice of each run, from which the median across slices was chosen. The median number of principal components ranged from 4–14 across each of the three datasets evaluated here, save for a couple of large outliers in the resting state data **(Fig. 2A)**. When the slice-wise parallel analysis output was considered, particularly in the hand-grasp task data, higher variability across subjects was observed in the inferior slices, which may be associated with lower signal quality in these regions (Hemmerling et al., 2023a), or different noise characteristics for this task. Lower signal quality may be attributed to the distance of inferior segments from the coil, or the documented time-varying respiration-induced differences in the B_0_ field that are higher at the lower cervical vertebrae, near the rostral part of the lungs (Vannesjo et al., 2018; Verma & Cohen-Adad, 2014). This variability is averaged out when selecting the median number of principal components across slices for each fMRI dataset.

The parallel analysis is computationally intensive, taking us well over an hour just to define the number of regressors to include. (Note, this analysis was done on a high-performance computing cluster and may not reflect timing for all computer systems/datasets.) However, the consistency of the cutoff number of principal components across the three datasets also implies that an appropriate number of components for a study could be reasonably extrapolated from a parallel analysis performed on a randomly selected subset of subjects, rather than on every run in the dataset, thus reducing the overall computational burden. Additionally, there is variability in the cutoff number of principal components across individual datasets, with low repeatability between Scan 1 and Scan 2 for the hand-grasping task dataset (ICC=0.316) **(Fig. S13)** and moderate repeatability for the breath-holding task dataset (ICC=0.591). This variability between repeated scans may be influenced by differences in motion or natural physiological fluctuations, as these scans were acquired a few hours apart. However, the median across subjects is similar, so performing the parallel analysis on a randomly selected subset of subjects may still be a reasonable option to reduce computational intensity. It is worth pointing out here that the number of components indicated by the parallel analysis appears to be affected by the number of timepoints (see **Fig. S3**) and may relate to the overall motion present in the data (see **Fig. S4**), so the results presented in **Fig. 2** should not be extrapolated for other datasets.

The two resting state B scans that had a large cutoff number of principal components also had some of the lowest motion in the resting state dataset. This may initially raise concerns, as we are suggesting that SpinalCompCor should include the greatest number of nuisance regressors when there is the smallest amount of anticipated motion in the data. However, in this case the noise may be arising from more diverse sources with more spatially varying temporal signatures, spreading it across a greater number of components. If a subject has a large movement artifact (i.e., bulk motion affecting the entire slice or volume), it can supersede these more subtle and local physiological fluctuations across the entire field of view. Indeed, the positive relationship between the first eigenvalue and motion supports this explanation, whereby data with larger motion-related artifacts have more variance explained by the first component.

### 4.3 Modeling with PCA-derived SpinalCompCor regressors

In the Extended model, SpinalCompCor noise regressors are added to our existing denoising model that already contains motion and physiological noise regressors, rather than directly comparing the two as separate models. This allows us to investigate if adding these extra data-driven regressors is useful *beyond* the denoising we are already doing as is standard in the field (**Table S1**; (Braaß et al., 2023; Dabbagh et al., 2024; Kinany et al., 2023; Kowalczyk et al., 2024; Landelle et al., 2023; Seifert et al., 2024)). These models typically include RETROICOR, motion, and CSF regressors, albeit with variation in the number and the specific methodology to calculate the regressors.

In the SpinalCompCor model, the PCA regressors are used as a replacement for physiological recording-derived RETROICOR regressors. This model allows us to investigate whether the SpinalCompCor method may be used as a suitable replacement for RETROICOR regressors. Future work evaluating a wider variety of models may reduce the overall number of nuisance regressors, thus maximizing the degrees of freedom in the GLM. However, typical spinal cord fMRI datasets are long, with over a hundred timepoints, suggesting that the addition of 4-14 SpinalCompCor regressors may not be overly problematic.

#### 4.3.1 Spatial variation of SpinalCompCor regressor parameter estimates

Before evaluating model performance, we probed what the origin of the SpinalCompCor regressors might be. The SpinalCompCor method relies on the assumption that structured noise sources outside the spinal cord impart or concur with time-varying noise within spinal cord tissue, such that this variance within the cord can be explained by regressors derived from the noise ROI outside the cord. The anatomical structures around the cervical spinal cord are primarily CSF in the subarachnoid space, blood vessels, the trachea, vertebrae, and neck muscles (Brooks et al., 2008). Noise would primarily be expected to be imparted from cardiac (e.g., CSF, vessels) and respiratory (e.g., trachea) sources (Eippert et al., 2017), with quasi-periodic time-varying cycles, or from motion.

The parameter estimate maps associated with the SpinalCompCor regressors tell us where these regressors are fitting noise *outside* of the spinal cord region, alluding to what noise sources the components might be representing. Parameter estimate maps from fMRI runs for which the GLM was rerun for the entire field of view, show interesting spatial information about the principal components. (As there are many principal component parameter estimate maps, we selected illustrative examples of the anatomical structure in the maps but also acknowledge that these examples are not fully representative.)

Greater parameter estimates are frequently observed in CSF and blood vessels – structures affected by the cardiac cycle **(e.g., Fig. 3A,B)**. **Figure S5** shows that white matter voxels adjacent to CSF may occasionally contain elevated parameter estimates, even when primarily within CSF voxels. This indicates some partial volume effects between the CSF and white matter or potential confounding effects of the pulsatile CSF on white matter edge voxels. As noted previously, there is already a separate slice-wise CSF regressor in the model, which is the average timeseries of the CSF voxels with the top 20% variance (Brooks et al., 2008; Kong et al., 2012). This established approach for extracting a nuisance regressor from CSF voxels may mask pulsatile effects that vary (i.e., “travel”) along the length of the cord (Hemmerling et al., 2022), which SpinalCompCor regressors can potentially address. Another area showing greater parameter estimates is in the neck muscles **(e.g., Fig. 3D)**. This may be related to respiration, as Brooks et al. (2008) observed respiratory effects in these tissues, but may also reflect any number of other physiological fluctuations occurring during scanning, such as swallowing, which we are unable to specifically observe in these data. Future work could integrate a periodic swallowing task, specifically aimed to isolate and quantify the impact of swallowing on spinal cord fMRI timeseries. Elevated parameter estimates at the edges of structures across the field of view may represent cases for which the SpinalCompCor regressor is explaining bulk motion **(e.g., Fig. S6C)**. Most maps appear to have some structure; however, a more unstructured example is seen in **Fig. 3C**.

A wider FOV in the resting state B dataset commonly shows higher parameter estimates in voxels that might be influenced by swallowing or contain air-filled respiratory structures **(Fig. S8)**. While the noise ROI does not actually cover this more ventral part of the image, the parameter estimate maps suggest that signal fluctuation in this region may be influential enough to affect voxels in the nearby noise ROI. This highlights the utility of acquisitions that limit the impact of noise confounds from outside of the imaged region, such as a reduced FOV acquisition with outer volume suppression (OVS) or inner FOV imaging with selective excitation (Kinany et al., 2022). The latter of these was used for the breath-hold task, hand-grasp task, and resting state A datasets in this study (specifically, ZOOMit from Siemens). Kinany et al. (2022) specifically compares OVS and ZOOMit for detection of task and resting state fMRI activity, demonstrating strengths and weaknesses of the approaches and their ability to image the spinal cord.

#### 4.3.2 Model evaluation

Next, we aimed to test whether SpinalCompCor regressors, reflecting structured noise from outside of the spinal cord, explained significant signal variance *inside* of the spinal cord. We performed a voxel-wise *F*-test to evaluate whether the Extended (i.e., full) model fit the hand-grasp task fMRI data significantly better than the Base (i.e., reduced) model. All spinal cord segments do not benefit similarly, as supported by the mixed modeling approach looking at the proportion of statistically significant voxels (*F*-test) across spinal cord segments **(Fig. 4)**. It is possible there are differences in the noise profile rostrocaudally along the cord contributing to the differential benefit from modeling with SpinalCompCor regressors. It would be interesting to determine whether this variability in SpinalCompCor benefit would persist for fMRI data in other regions such as the lumbar cord.

Most breath-hold and hand-grasp fMRI runs saw SpinalCompCor explain significantly more variance in 10–60% of voxels in both white and gray matter regions (based on the *F*-test). However, there is substantial variability in this percentage, suggesting SpinalCompCor may have a variable impact on modeling across subjects and runs. Extracting these tissue-specific metrics requires warping individual functional datasets to the PAM50 template, so these percentage metrics will inevitably capture some partial volume and misregistration effects that may contribute to this variability. Although the masks were visually checked during preprocessing, the imprecision in tissue type delineation at this field strength and spatial resolution is important to recognize. Still, these results indicate that the SpinalCompCor regressors, derived from a noise ROI outside of the spinal cord, do indeed explain significant additional variance within the spinal cord tissues, beyond variance explained by a standard set of motion and physiological noise regressors in our fMRI data.

Notably, most resting state A fMRI runs saw SpinalCompCor explain significant additional variance in less than 25% of voxels across regions, compared to 90-100% of voxels for most resting state B fMRI runs. This suggests that differences in acquisition may be the most influential driver of the performance of SpinalCompCor and similar methods. Specifically, the SpinalCompCor method appears, at the subject-level, to be more beneficial for the 3D acquisition as compared to the 2D inner FOV acquisition.

#### 4.3.3 Effect on downstream group-level activation results

We evaluated the utility of SpinalCompCor in improving sensitivity and specificity to motor neural activity and vascular reactivity using the hand-grasping task and breath-holding task fMRI datasets, respectively. The Base model included a base set of nuisance regressors (see section 2.2.4), and the Extended model additionally included the number of component regressors indicated by the parallel analysis for the full run, ranging from 7–19 (hand-grasp) and 6-16 (breath-hold) across the dataset studied (two runs/subject). Hand-grasping motor activation is primarily expected to localize (within the imaging field of view) to gray matter voxels in the C7-C8 spinal cord segments, ipsilateral to the side of the task (Kuypers, 1982; Leinberry & Wehbé, 2004). (See Hemmerling et al. (2023a) for a more detailed discussion of expected activity localization.)

Group-level modeling of hand-grasping task activation with SpinalCompCor regressors produced mixed results. For our lateralized hand-grasping task, we would primarily expect ipsilateral gray matter activation associated with motor activity (i.e., ventral horns) as well as some proprioceptive or sensory activity related to the task (i.e., dorsal horns). We would expect less activation in white matter voxels, given functional neurovascular anatomy and BOLD sensitivity at 3 T. Simplistically, when comparing the Base model to the Extended or SpinalCompCor model, a reduction in statistically significant activation in white matter voxels might be expected, whilst we might anticipate gray matter activation to stay the same or increase.

In white matter voxels, the proportion of active voxel changes to now active or no longer active is inconsistent between left and right grasping, suggesting that the benefit indicated at the individual dataset level **(Fig. 4)** does not translate to a consistent change in (potentially spurious) task activation in white matter regions. However, we note that white matter activation may not be entirely spurious; there is increasing evidence of meaningful neural BOLD signals in white matter in the brain (Gore et al., 2019), as well as in recent work in the primate spinal cord (Sengupta et al., 2023). Therefore, we cannot discount the white matter BOLD signal as containing true activation.

For right hand-grasping, in ventral gray matter in the C8 spinal cord segment – part of the hypothesized region in which peak motor activity would be expected (Vanderah & Gould, 2016) – the proportion of voxels that are now active vs. no longer active with the SpinalCompCor or Extended model is larger, suggesting our method improved sensitivity to true neural activation in the BOLD timeseries. However, in the corresponding ROI for left hand-grasping, the proportion of voxels that are now active with the SpinalCompCor or Extended model is smaller compared to those that are no longer active. In fact, in most ventral gray matter ROIs, a higher proportion of voxels are no longer active with the SpinalCompCor model, while the opposite is seen for many dorsal gray matter ROIs. Without having data on hand dominance or a graded force-level motor task, we can speculate that differences in hand dominance and the absolute force being generated by each hand may lead to differences in absolute neural activity as well as correlated noise artifacts in left versus right hand-grasping. These differences may also cause variation in the sensory activity associated with the motor task, impacting our dorsal horn results. Alternatively, we previously observed differences in the strength and timing of the vascular response between ventral and dorsal regions in the cord – specifically, we observed a stronger and earlier ventral response compared to the dorsal response (Hemmerling et al., 2025). These vascular differences may also impact the observed motor activity and change in motor activity when using SpinalCompCor, which also shows differences between ventral and dorsal regions. Finally, it is also possible that SpinalCompCor is spuriously removing or adding active voxels in these regions, rather than effectively addressing noise. Regardless of the reasons, our observations suggest that SpinalCompCor is impacting task activation mapping in an unpredictable way. Moreover, there may be collinearity between SpinalCompCor regressors and our task regressors that must be addressed to ensure SpinalCompCor is appropriate in task fMRI datasets.

Breath-holding-induced hypercapnia is a systemic vasodilatory stimulus. Active voxels would be expected across spinal cord tissue; however, based on our previous work, without accounting for temporal differences in the local response, voxel activation would be most expected in ventral gray matter (Hemmerling et al., 2025). Changes in active voxels between models were primarily observed at the edges of clusters of active voxels. In the ventral gray matter, about 7-19% of voxels are no longer active between the models. These changes could be due to partial volume effects or decreased sensitivity at these boundary regions. Across both the hand-grasping and breath-holding tasks, the difference between the Base-Extended and Base-SpinalCompCor models appear very similar. Importantly, because breath-holding evokes a systemic vasodilatory response, there may be increased common signal fluctuations both inside and outside of the spinal cord. In accounting for this, SpinalCompCor may inadvertently regress out this shared, task-related variance, thereby reducing parameter estimates for the task (CO_2_) regressor and the percent significant voxels.

#### 4.3.4 Effect on resting state functional connectivity

The utility of SpinalCompCor for denoising resting state fMRI data and improving functional connectivity estimates was also evaluated for both resting state datasets. We may expect established connections, such as dorsal-dorsal or ventral-ventral (Barry et al., 2014), to strengthen with improved denoising. However, this was not seen for either of the resting state datasets. In the resting state B dataset, the Base model instead has a significantly higher correlation compared to the Extended and SpinalCompCor models for all comparisons.

Considering the high proportion of significant voxels from the F-test for this dataset, it is highly likely this represents the removal of spurious noise-driven correlations when using SpinalCompCor regressors. However, we cannot discount the possibility that there is correlation between intrinsic resting state fluctuations with signals outside of the spinal cord, and that this reduction represents the removal of real neural connectivity between regions. The difference in acquisition type and parameters may be driving the difference in functional connectivity between the two resting state datasets. Future research would be necessary to understand the exact impacts to resting state fMRI.

#### 4.3.5 Collinearity

The inconsistent hand-grasp task activation mapping results led us to consider whether there could be any issues with regressor collinearity interfering with our statistical inference (e.g., cross-talk between the SpinalCompCor nuisance regressors and our task regressors of interest). In particular, when including additional regressors, it would be cause for concern if there is a strong correlation between one of these and the task regressor(s). A correlation matrix of model regressors for an example hand-grasp task model (Subject 13, Scan 1, Slice 12) shows the collinearity of each regressor with the others **(Fig. 10A,B)**. In this example, high correlations are observed between nuisance regressors with other nuisance regressors, which indicates some redundancy within our model. In fact, this provides additional insight into what motion or physiological confounds SpinalCompCor is capturing. For example, PC1 and PC2 correlate strongly with CSF and cardiac RETROICOR regressors, respectively. Thus, in the absence of external physiological recordings, using data-driven PCA regressors like these may be particularly beneficial. However, as shown by the spatial correlation **(Fig. 5),** the SpinalCompCor regressors are likely not a perfect substitute for the RETROICOR regressors which require high-quality physiological recordings.

**Fig. 10.**
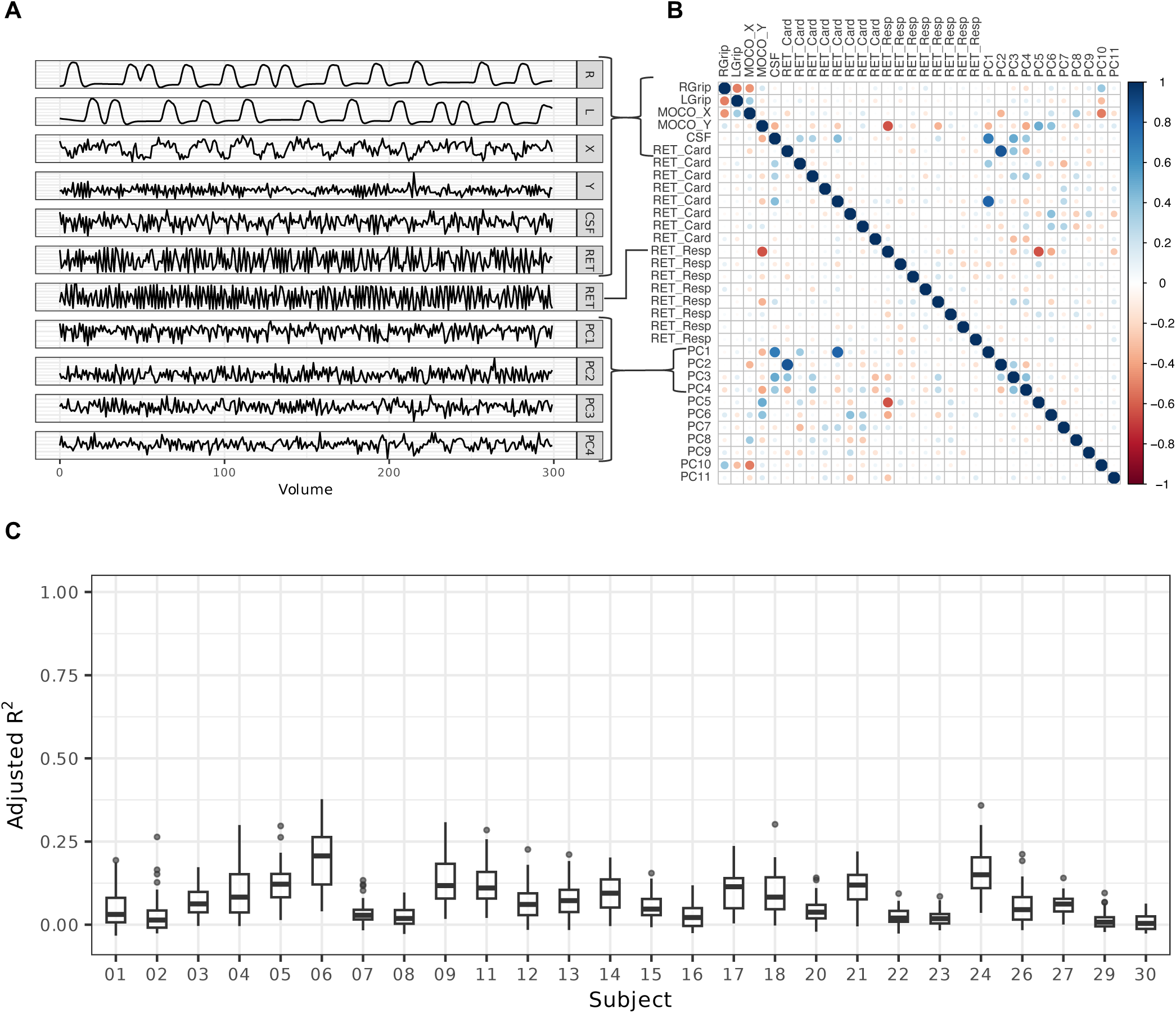
Collinearity of fMRI models for an example run and across all hand-grasp data. **(A)** Select regressors for example Extended model (Subject 13, Scan 1, Slice 12). **(B)** Example correlation matrix for the same model. **(C)** Adjusted R^2^ for the model of Task ∼ PC1 + PC2 + …+ PCn across all slices, subjects, and scans.

Interestingly, there are some non-zero correlations indicated between principal component regressors when they should be orthogonal to each other (lower right quadrant of **Fig. 10B**). This is due to FSL’s modeling (e.g., FEAT was set up to apply a high-pass filter, which is applied to both the fMRI timeseries and the regressors). In other words, the principal components initially had zero correlation, but the regressors that were actually used in the model, and evaluated here, regained some small degree of correlation following temporal filtering.

Because our chief interest is whether the principal component regressors would share variance with the task regressors, and thus negatively impact the parameter estimates/inferences for task activation, linear models of the task vs. principal component regressors (Task ∼ PC1 + PC2 + … + PCn) were computed and the associated adjusted R^2^ was plotted for every slice of each fMRI run **(Fig. 10C)**. Most adjusted *R^2^* values are low, so we are not particularly concerned with introducing model collinearity when adding principal component regressors. However, there some outliers with higher correlations which could contribute to the ambiguous group-level activation results.

### 4.4 Recommendations for use

The choice of whether to use SpinalCompCor for deriving nuisance regressors is nuanced. A summary of the data included in this study is provided for reference **(Table 3)**. The effect of SpinalCompCor on these datasets was varied; based on overall findings, we lay out here some guidelines for use.

**Table 3.** Summary of data acquisitions and model regressors.

| Dataset | Acquisition Type | Volumes | Number of Nuisance Regressors for Modeling |  |  |  |  |
| --- | --- | --- | --- | --- | --- | --- | --- |
|  |  |  | SpinalCompCor<br>Mean (SD)<br>*Models 2,3* | Motion<br>2 DOF<br>*Models 1,2,3* | CSF<br>*Models 1,2,3* | Cardiac<br>RETROICOR<br>*Models 1,2* | Respiratory<br>RETROICOR<br>*Models 1,2* |
| Breath-Hold Task | 2D, gradient-echo EPI, inner FOV (ZOOMit), Siemens | 205 | 10.3 (1.79) | 2 | 1 | 8 | 8 |
| Hand-Grasp Task | 2D, gradient-echo EPI, inner FOV (ZOOMit), Siemens | 300 | 15.2 (2.93) | 2 | 1 | 8 | 8 |
| Resting State A | 2D, gradient-echo EPI, inner FOV (ZOOMit), Siemens | 160 | 9.83 (5.58) | 2 | 1 | 8 | 8 |
| Resting State B | 3D, gradient-echo multi-shot, Philips | 434 | 28.3 (23.3) | 2 | 1 | 8 | 8 |

Acquisition type, number of volumes, and number of nuisance regressors for modeling across each included dataset. For the SpinalCompCor regressor, the mean and standard deviation (SD) are reported. Whether each nuisance regressor type was included in the Base (1), Extended (2), or SpinalCompCor (3) model is indicated by the corresponding italicized numbers below each column header (see also section 2.2.4). DOF=degrees of freedom

Use of SpinalCompCor is recommended:

- In lieu of physiological recordings, whether they are poor quality or unavailable.
- With the aim of reducing false positives in task fMRI or inflated connectivity in resting-state fMRI.
- If the available degrees of freedom are sufficient.
- When the acquisition field of view is larger (e.g., non-inner FOV acquisition), encompassing more potential sources of physiological noise.

Use of SpinalCompCor is *not* recommended:

- If the available degrees of freedom are insufficient.
- When there is suspected correlation between noise fluctuations and the signal of interest inside the cord (e.g., task correlated motion, systemic vasodilation associated with a breath-hold task).

Ultimately, individual judgment should be used on specific datasets to determine suitability for this technique. In our data, SpinalCompCor (only) is best viewed as a practical fallback when physiological recordings are unavailable, rather than as a drop-in replacement for recording-based physiological regressors, as the similarity to recording-based denoising varied across datasets/acquisitions. It is also important to remember that results reported here, namely, the number of SpinalCompCor regressors, do not generalize to other acquisitions.

### 4.5 Limitations and future directions

Spinal cord fMRI data analysis is actively being improved upon by researchers in the community. As such, there were some small changes to data processing and analysis that we have made since the original work using the hand-grasp task dataset (Hemmerling et al., 2023a). While the original work did not observe many significant voxels at the C8 spinal cord segment, we have slightly improved the co-registration and are using the updated Spinal Cord Toolbox version (6.1.0) of the spinal cord segment template. Thus, increased activation in the C8 segment is observed here.

A main limitation of this work is that we selected and tested a specific analysis pipeline and specific collection of nuisance regressors in our Base model. While the basic fMRI processing steps used are standard, they may be implemented differently by different research groups, depending both on software choice and individuals’ use of the tools. The selection of regressors to include in the GLM for the Base model is based on published literature **(Table S1)**, the availability of physiological data recordings, and what has been successful in our own past work.

This work incorporated four 3 T fMRI task and resting state datasets collected on scanners from two different vendors. This variation should increase the applicability of the results of this work. However, the full gamut of possible vendors, scan parameters, tasks, et cetera, cannot be represented by these datasets alone. Furthermore, it seems that the acquisition type and parameters may be more influential than initially anticipated, indicating that it may be important to tailor denoising strategies for specific acquisitions. The parallel analysis presented here is an automated and standardized strategy for determining how to appropriately implement SpinalCompCor across varied datasets.

There is a growing initiative for sharing spinal cord fMRI data across the international research community (Banerjee et al., 2025) and such resources will be invaluable for further standardizing methods in the spinal cord fMRI field, just as big data initiatives in brain fMRI like the Human Connectome Project have done in the last two decades (Elam et al., 2021).

Each of these choices affects the results and interpretation of the efficacy of using PCA-derived SpinalCompCor regressors in spinal cord fMRI modeling. Ultimately, we observe that SpinalCompCor can be applied in a systematic, principled manner, and explains significant variance within the spinal cord parenchyma in a variety of datasets. However, the impact of such denoising on task activation and functional connectivity was ambiguous. In the hand-grasping task data, the effects may be related to task correlated variance in the SpinalCompCor regressors for the hand-grasping task. The nuisance regression aims to remove motion and physiological effects, however, regressors exhibiting task-correlated fluctuations could unintentionally reduce the share of variance that task regressors receive, hindering the detection of activation. (Note that such collinearity between SpinalCompCor regressors and spontaneous signal fluctuations in the resting state data would present a similar challenge, although without ground truth knowledge of the intrinsic neurovascular signals in the spinal cord it is even more difficult to tease these effects apart.) Further development of this method could take inspiration from literature in the following ways. In the CompCor method, Behzadi et al. (2007) described a technique to remove stimulus-related components: voxels are excluded from the noise ROI that are too correlated with the stimulus-related reference function. A similar technique could be tested for spinal cord noise ROIs. Alternatively, PCA-derived SpinalCompCor regressors could be orthogonalized with respect to other regressors in the model. However, this should be implemented with caution, and it is advisable to fully understand warnings related to orthogonalization in fMRI modeling (see Mumford et al. (2015)).

A different technique that might help to address task-correlated noise is the use of multi-echo independent component analysis (Kundu et al., 2012). Previous work has shown this method to be beneficial in the presence of high task-correlated head motion at the subject-level (Reddy et al., 2024). However, group-level results were similar across levels of head motion (Reddy et al., 2024), indicating that group-level activation mapping may already be fairly robust to variation in noise across subjects. This is similar to what was observed here in the hand-grasp task fMRI data – the Extended model explained noise at the subject-level, but a corresponding change in the direction of hypothesized motor activation for group-level maps was not seen. Therefore, advanced denoising methods like SpinalCompCor may be particularly beneficial for precision mapping of the individual spinal cord.

## 5 CONCLUSIONS

In this work, we presented a method to derive PCA-based nuisance regressors from a noise ROI for spinal cord fMRI modeling. Components were designated as regressors for denoising using a parallel analysis comparing real and simulated noise ROI timeseries. This automated and principled approach suggested a reasonable number of components to retain as regressors across four diverse datasets. Across datasets, SpinalCompCor regressors reflected variance in areas of the fMRI volume *outside* the spinal cord that correspond to anatomical features expected to exhibit physiological fluctuations and systemically confound fMRI timeseries (e.g., blood vessels, CSF). Accordingly, these regressors explained significant signal variance *inside* the spinal cord in subject-level modeling. The difference in subject-level results for the two resting state fMRI datasets, acquired using different scan acquisition protocols on different scanners, shows that the effectiveness of the denoising strategy may be primarily dependent on details of the fMRI acquisition, more than the presence or absence of a task paradigm. Mixed results were observed when considering the effects of SpinalCompCor on group-level task activation mapping and functional connectivity analysis, and this may suggest challenges with task-correlated noise and model collinearity. Overall, we demonstrate the benefits of using PCA-derived SpinalCompCor regressors in modeling spinal cord fMRI timeseries. However, we show that these benefits may not be equal across fMRI datasets and may be acquisition dependent.

## Data and Code Availability

Code to create regressors and perform the parallel analysis is available at https://github.com/BrightLab-ANVIL/spinalcompcor. Datasets from three sources were used and are available according to their original publication data availability. The hand-grasping task fMRI data are publicly available on OpenNeuro at doi:10.18112/openneuro.ds004616.v1.1.1 (Hemmerling et al., 2023b). The breath-holding task fMRI data are publicly available on OpenNeuro at doi:10.18112/openneuro.ds004974.v1.0.0 (Hemmerling et al., 2024b). The resting state B fMRI data are available upon reasonable request (see Barry et al. (2018)).

## Author Contributions

K.J.H: Conceptualization, Methodology, Software, Formal analysis, Investigation, Data curation, Writing - Original Draft, Writing - Review & Editing, Visualization, Project administration. A.D.V.: Methodology, Writing - Review & Editing. C.G.: Software, Formal analysis, Writing - Review & Editing. R.L.B.: Conceptualization, Methodology, Investigation, Data curation, Writing - Review & Editing, Funding acquisition. M.G.B.: Conceptualization, Methodology, Writing - Review & Editing, Supervision, Project administration, Funding acquisition

## Acknowledgments

This work was supported by the Center for Translational Imaging at Northwestern University. This research was also supported in part through the computational resources and staff contributions provided for the Quest high performance computing facility at Northwestern University which is jointly supported by the Office of the Provost, the Office for Research, and Northwestern University Information Technology. The authors are grateful to Dr. Kenneth A. Weber II for generously sharing a resting state fMRI dataset (“A”) used in our analysis.

## Footnotes

1 The IAAFT algorithm described here was based on a webpage that is no longer accessible. The cited work is from the same author; the IAAFT description can be found in Venema et al. 2006 (section 2).

## References

Banerjee, R., Kaptan, M., Tinnermann, A., Khatibi, A., Dabbagh, A., Büchel, C., Kündig, C. W., Law, C. S. W., Pfyffer, D., Lythgoe, D. J., Tsivaka, D., Van De Ville, D., Eippert, F., Muhammad, F., Glover, G. H., David, G., Haynes, G., Haaker, J., Brooks, J. C. W., … Cohen-Adad, J. (2025). EPISeg: Automated segmentation of the spinal cord on echo planar images using open-access multi-center data. Imaging Neuroscience, 3. 10.1162/IMAG.a.98

Barry, R. L., Conrad, B. N., Smith, S. A., & Gore, J. C. (2018). A practical protocol for measurements of spinal cord functional connectivity. Scientific Reports, 8(1), 1–10. 10.1038/s41598-018-34841-6

Barry, R. L., Rogers, B. P., Conrad, B. N., Smith, S. A., & Gore, J. C. (2016). Reproducibility of resting state spinal cord networks in healthy volunteers at 7 Tesla. NeuroImage, 133, 31–40. 10.1016/j.neuroimage.2016.02.058

Barry, R. L., Smith, S. A., Dula, A. N., & Gore, J. C. (2014). Resting state functional connectivity in the human spinal cord. ELife, 2014(3). 10.7554/eLife.02812

Behzadi, Y., Restom, K., Liau, J., & Liu, T. T. (2007). A component based noise correction method (CompCor) for BOLD and perfusion based fMRI. Human Brain Mapping Journal, 37(1), 90–101. 10.1016/j.neuroimage.2007.04.042

Braaß, H., Feldheim, J., Chu, Y., Tinnermann, A., Finsterbusch, J., Büchel, C., Schulz, R., & Gerloff, C. (2023). Association between activity in the ventral premotor cortex and spinal cord activation during force generation—A combined cortico-spinal fMRI study. Human Brain Mapping, 44(18). 10.1002/hbm.26523

Brooks, J. C. W., Beckmann, C. F., Miller, K. L., Wise, R. G., Porro, C. A., Tracey, I., & Jenkinson, M. (2008). Physiological noise modelling for spinal functional magnetic resonance imaging studies. NeuroImage, 39(2), 680–692. 10.1016/j.neuroimage.2007.09.018

Caballero-Gaudes, C., & Reynolds, R. C. (2017). Methods for cleaning the BOLD fMRI signal. NeuroImage, 154. 10.1016/j.neuroimage.2016.12.018

Combes, A., Narisetti, L., Sengupta, A., Rogers, B. P., Sweeney, G., Prock, L., Houston, D., McKnight, C. D., Gore, J. C., Smith, S. A., & O’Grady, K. P. (2023). Detection of resting-state functional connectivity in the lumbar spinal cord with 3T MRI. Scientific Reports, 13(1), 18189. 10.1038/s41598-023-45302-0

Dabbagh, A., Horn, U., Kaptan, M., Mildner, T., Müller, R., Lepsien, J., Weiskopf, N., Brooks, J. C. W., Finsterbusch, J., & Eippert, F. (2024). Reliability of task-based fMRI in the dorsal horn of the human spinal cord. Imaging Neuroscience, 2, 1–27. 10.1162/IMAG_A_00273

De Leener, B., Fonov, V. S., Collins, D. L., Callot, V., Stikov, N., & Cohen-Adad, J. (2018). PAM50: Unbiased multimodal template of the brainstem and spinal cord aligned with the ICBM152 space. NeuroImage, 165, 170–179. 10.1016/j.neuroimage.2017.10.041

Deshpande, R., & Barry, R. (2022). Neptune: a toolbox for spinal cord functional MRI data processing and quality assurance. Proc Intl Soc Mag Reson Med 30. 10.58530/2022/0396

Eippert, F., Kong, Y., Jenkinson, M., Tracey, I., & Brooks, J. C. W. (2017). Denoising spinal cord fMRI data: Approaches to acquisition and analysis. NeuroImage, 154, 255–266. 10.1016/j.neuroimage.2016.09.065

Elam, J. S., Glasser, M. F., Harms, M. P., Sotiropoulos, S. N., Andersson, J. L. R., Burgess, G. C., Curtiss, S. W., Oostenveld, R., Larson-Prior, L. J., Schoffelen, J. M., Hodge, M. R., Cler, E. A., Marcus, D. M., Barch, D. M., Yacoub, E., Smith, S. M., Ugurbil, K., & Van Essen, D. C. (2021). The Human Connectome Project: A retrospective. NeuroImage, 244. 10.1016/j.neuroimage.2021.118543

Glover, G. H., Li, T. Q., & Ress, D. (2000). Image-based method for retrospective correction of physiological motion effects in fMRI: RETROICOR. Magnetic Resonance in Medicine, 44(1), 162–167. 10.1002/1522-2594(200007)44:1<162::AID-MRM23>3.0.CO;2-E

Gore, J. C., Li, M., Gao, Y., Wu, T. L., Schilling, K. G., Huang, Y., Mishra, A., Newton, A. T., Rogers, B. P., Chen, L. M., Anderson, A. W., & Ding, Z. (2019). Functional MRI and resting state connectivity in white matter - a mini-review. Magnetic Resonance Imaging, 63. 10.1016/j.mri.2019.07.017

Gros, C., De Leener, B., Badji, A., Maranzano, J., Eden, D., Dupont, S. M., Talbott, J., Zhuoquiong, R., Liu, Y., Granberg, T., Ouellette, R., Tachibana, Y., Hori, M., Kamiya, K., Chougar, L., Stawiarz, L., Hillert, J., Bannier, E., Kerbrat, A., … Cohen-Adad, J. (2019). Automatic segmentation of the spinal cord and intramedullary multiple sclerosis lesions with convolutional neural networks. NeuroImage, 184, 901–915. 10.1016/J.NEUROIMAGE.2018.09.081

Hemmerling, K. J., Hoggarth, M. A., Parrish, T., & Bright, M. G. (2022). Spinal cord fMRI heatmaps reveal a structured cardiac artifact “traveling” along the length of the cord. Proceedings 30th Scientific Meeting, International Society for Magnetic Resonance in Medicine.

Hemmerling, K. J., Hoggarth, M. A., Sandhu, M. S., Parrish, T. B., & Bright, M. G. (2023a). Spatial distribution of hand-grasp motor task activity in spinal cord functional magnetic resonance imaging. Human Brain Mapping, 44(17), 5567–5581. 10.1002/hbm.26458

Hemmerling, K. J., Hoggarth, M. A., Sandhu, M. S., Parrish, T. B., & Bright, M. G. (2023b). Spatial distribution of hand-grasp motor task activity in spinal cord functional magnetic resonance imaging. OpenNeuro.

Hemmerling, K. J., Hoggarth, M. A., Sandhu, M. S., Parrish, T. B., & Bright, M. G. (2024). MRI mapping of hemodynamics in the human spinal cord. OpenNeuro.

Hemmerling, K. J., Hoggarth, M. A., Sandhu, M. S., Parrish, T. B., & Bright, M. G. (2025). MRI mapping of hemodynamics in the human spinal cord. Scientific Reports, 15(1), 34880. 10.1038/s41598-025-17048-4

Hu, Y., Jin, R., Li, G., Luk, K. D. K., & Wu, E. X. (2018). Robust spinal cord resting-state fMRI using independent component analysis-based nuisance regression noise reduction. Journal of Magnetic Resonance Imaging, 48(5), 1421–1431. 10.1002/jmri.26048

Jenkinson, M., Beckmann, C. F., Behrens, T. E. J., Woolrich, M. W., & Smith, S. M. (2012). FSL. NeuroImage, 62(2), 782–790. 10.1016/j.neuroimage.2011.09.015

Kaptan, M., Horn, U., Vannesjo, S. J., Mildner, T., Weiskopf, N., Finsterbusch, J., Brooks, J. C. W., & Eippert, F. (2023). Reliability of resting-state functional connectivity in the human spinal cord: Assessing the impact of distinct noise sources. NeuroImage, 275. 10.1016/j.neuroimage.2023.120152

Kinany, N., Khatibi, A., Lungu, O., Finsterbusch, J., Büchel, C., Marchand-Pauvert, V., Van De Ville, D., Vahdat, S., & Doyon, J. (2023). Decoding cerebro-spinal signatures of human behavior: Application to motor sequence learning. NeuroImage, 275. 10.1016/j.neuroimage.2023.120174

Kinany, N., Pirondini, E., Mattera, L., Martuzzi, R., Micera, S., & Van De Ville, D. (2022). Towards reliable spinal cord fMRI: Assessment of common imaging protocols. NeuroImage, 250. 10.1016/j.neuroimage.2022.118964

Kong, Y., Jenkinson, M., Andersson, J., Tracey, I., & Brooks, J. C. W. (2012). Assessment of physiological noise modelling methods for functional imaging of the spinal cord. NeuroImage, 60(2), 1538–1549. 10.1016/j.neuroimage.2011.11.077

Kowalczyk, O. S., Medina, S., Tsivaka, D., McMahon, S. B., Williams, S. C. R., Brooks, J. C. W., Lythgoe, D. J., & Howard, M. A. (2024). Spinal fMRI demonstrates segmental organisation of functionally connected networks in the cervical spinal cord: A test–retest reliability study. Human Brain Mapping, 45(2). 10.1002/hbm.26600

Kundu, P., Inati, S. J., Evans, J. W., Luh, W. M., & Bandettini, P. A. (2012). Differentiating BOLD and non-BOLD signals in fMRI time series using multi-echo EPI. NeuroImage, 60(3). 10.1016/j.neuroimage.2011.12.028

Kuypers, H. G. J. M. (1982). A New Look at the Organization of the Motor System. Progress in Brain Research, 57(C). 10.1016/S0079-6123(08)64138-2

Landelle, C., Dahlberg, L. S., Lungu, O., Misic, B., De Leener, B., & Doyon, J. (2023). Altered Spinal Cord Functional Connectivity Associated with Parkinson’s Disease Progression. Movement Disorders, 38(4). 10.1002/mds.29354

Leinberry, C. F., & Wehbé, M. A. (2004). Brachial plexus anatomy. Hand Clinics, 20(1). 10.1016/S0749-0712(03)00088-X

Lindquist, M. A., Geuter, S., Wager, T. D., & Caffo, B. S. (2019). Modular preprocessing pipelines can reintroduce artifacts into fMRI data. Human Brain Mapping, 40(8). 10.1002/hbm.24528

Mumford, J. A., Poline, J. B., & Poldrack, R. A. (2015). Orthogonalization of regressors in fMRI models. PLoS ONE, 10(4). 10.1371/journal.pone.0126255

Nichols, T. E., & Holmes, A. P. (2002). Nonparametric Permutation Tests For Functional Neuroimaging: A Primer with Examples. Human Brain Mapping, 15(1), 1–25. 10.1002/hbm.1058

Patriat, R., Molloy, E. K., & Birn, R. M. (2015). Using Edge Voxel Information to Improve Motion Regression for rs-fMRI Connectivity Studies. Brain Connectivity, 5(9), 582. 10.1089/BRAIN.2014.0321

Piché, M., Cohen-Adad, J., Nejad, M. K., Perlbarg, V., Xie, G., Beaudoin, G., Benali, H., & Rainville, P. (2009). Characterization of cardiac-related noise in fMRI of the cervical spinal cord. Magnetic Resonance Imaging, 27(3), 300–310. 10.1016/j.mri.2008.07.019

Raj, D., Anderson, A. W., & Gore, J. C. (2001). Respiratory effects in human functional magnetic resonance imaging due to bulk susceptibility changes. Physics in Medicine and Biology, 46(12), 3331–3340. 10.1088/0031-9155/46/12/318

Reddy, N. A., Zvolanek, K. M., Moia, S., Caballero-Gaudes, C., & Bright, M. G. (2024). Denoising task-correlated head motion from motor-task fMRI data with multi-echo ICA. Imaging Neuroscience, 2. 10.1162/imag_a_00057

Rieseberg, S., Frahm, J., & Finsterbusch, J. (2002). Two-dimensional spatially-selective RF excitation pulses in echo-planar imaging. Magnetic Resonance in Medicine, 47(6), 1186–1193. 10.1002/MRM.10157

Seifert, A. C., Xu, J., Kong, Y., Eippert, F., Miller, K. L., Tracey, I., & Vannesjo, S. J. (2024). Thermal stimulus task fMRI in the cervical spinal cord at 7 Tesla. Human Brain Mapping, 45(3). 10.1002/hbm.26597

Sengupta, A., Mishra, A., Wang, F., Chen, L., & Gore, J. (2023). Identification of synchronous BOLD signal patterns in white matter of primate spinal cord. Research Square. 10.21203/rs.3.rs-2389151/v1

Smith, S. M., & Nichols, T. E. (2009). Threshold-free cluster enhancement: Addressing problems of smoothing, threshold dependence and localisation in cluster inference. NeuroImage, 44(1), 83–98. 10.1016/J.NEUROIMAGE.2008.03.061

Vanderah, T. W., & Gould, D. J. (2016). Spinal Cord. In Nolte’s the Human Brain (7th ed., pp. 233–271). Elsevier Inc.

Vannesjo, S. J., Miller, K. L., Clare, S., & Tracey, I. (2018). Spatiotemporal characterization of breathing-induced B0 field fluctuations in the cervical spinal cord at 7T. NeuroImage, 167, 191–202. 10.1016/j.neuroimage.2017.11.031

Venema, V., Meyer, S., García, S. G., Kniffka, A., Simmer, C., Crewell, S., Löhnert, U., Trautmann, T., & Macke, A. (2006). Surrogate cloud fields generated with the iterative amplitude adapted Fourier transform algorithm. Tellus, Series A: Dynamic Meteorology and Oceanography, 58(1). 10.1111/j.1600-0870.2006.00160.x

Verma, T., & Cohen-Adad, J. (2014). Effect of respiration on the B0 field in the human spinal cord at 3T. Magnetic Resonance in Medicine, 72(6), 1629–1636. 10.1002/mrm.25075

Weber, K. A., Chen, Y., Wang, X., Kahnt, T., & Parrish, T. B. (2016a). Functional magnetic resonance imaging of the cervical spinal cord during thermal stimulation across consecutive runs. NeuroImage, 143. 10.1016/j.neuroimage.2016.09.015

Weber, K. A., Chen, Y., Wang, X., Kahnt, T., & Parrish, T. B. (2016b). Lateralization of cervical spinal cord activity during an isometric upper extremity motor task with functional magnetic resonance imaging. NeuroImage, 125, 233–243. 10.1016/j.neuroimage.2015.10.014

Weber, K. A., Sentis, A. I., Bernadel-Huey, O. N., Chen, Y., Wang, X., Parrish, T. B., & Mackey, S. (2018). Thermal Stimulation Alters Cervical Spinal Cord Functional Connectivity in Humans. Neuroscience, 369, 40–50. 10.1016/j.neuroscience.2017.10.035

Winkler, A. M., Ridgway, G. R., Webster, M. A., Smith, S. M., & Nichols, T. E. (2014). Permutation inference for the general linear model. NeuroImage, 92(100), 381–397. 10.1016/J.NEUROIMAGE.2014.01.060

Xie, G., Piché, M., Khoshnejad, M., Perlbarg, V., Chen, J. I., Hoge, R. D., Benali, H., Rossignol, S., Rainville, P., & Cohen-Adad, J. (2012). Reduction of physiological noise with independent component analysis improves the detection of nociceptive responses with fMRI of the human spinal cord. NeuroImage, 63(1), 245–252. 10.1016/j.neuroimage.2012.06.057

