## Supplemental Material for "Data-driven denoising in spinal cord fMRI with principal component analysis"

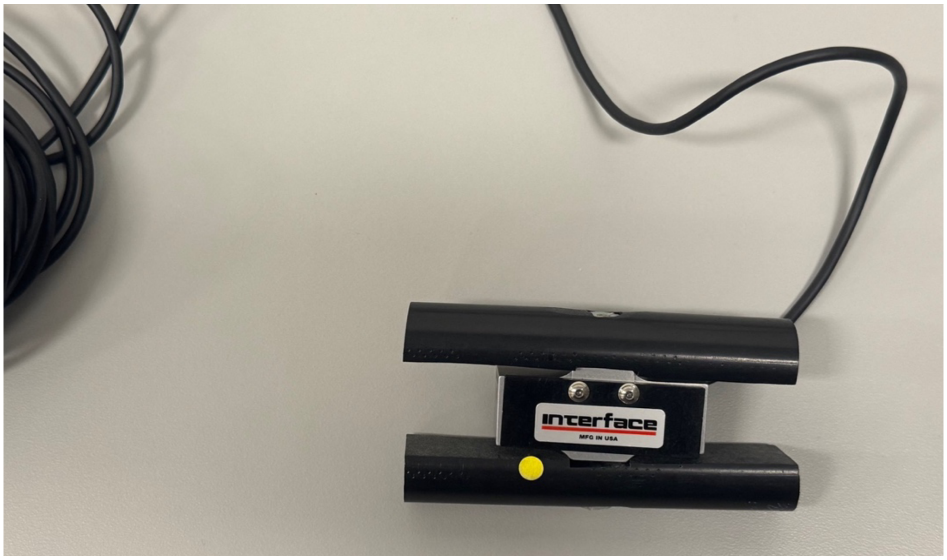


**Fig. S1. Custom MRI-safe hand grip.** One of two hand grips made from affixing two halves of a Delrin rod to an MRI compatible load cell.

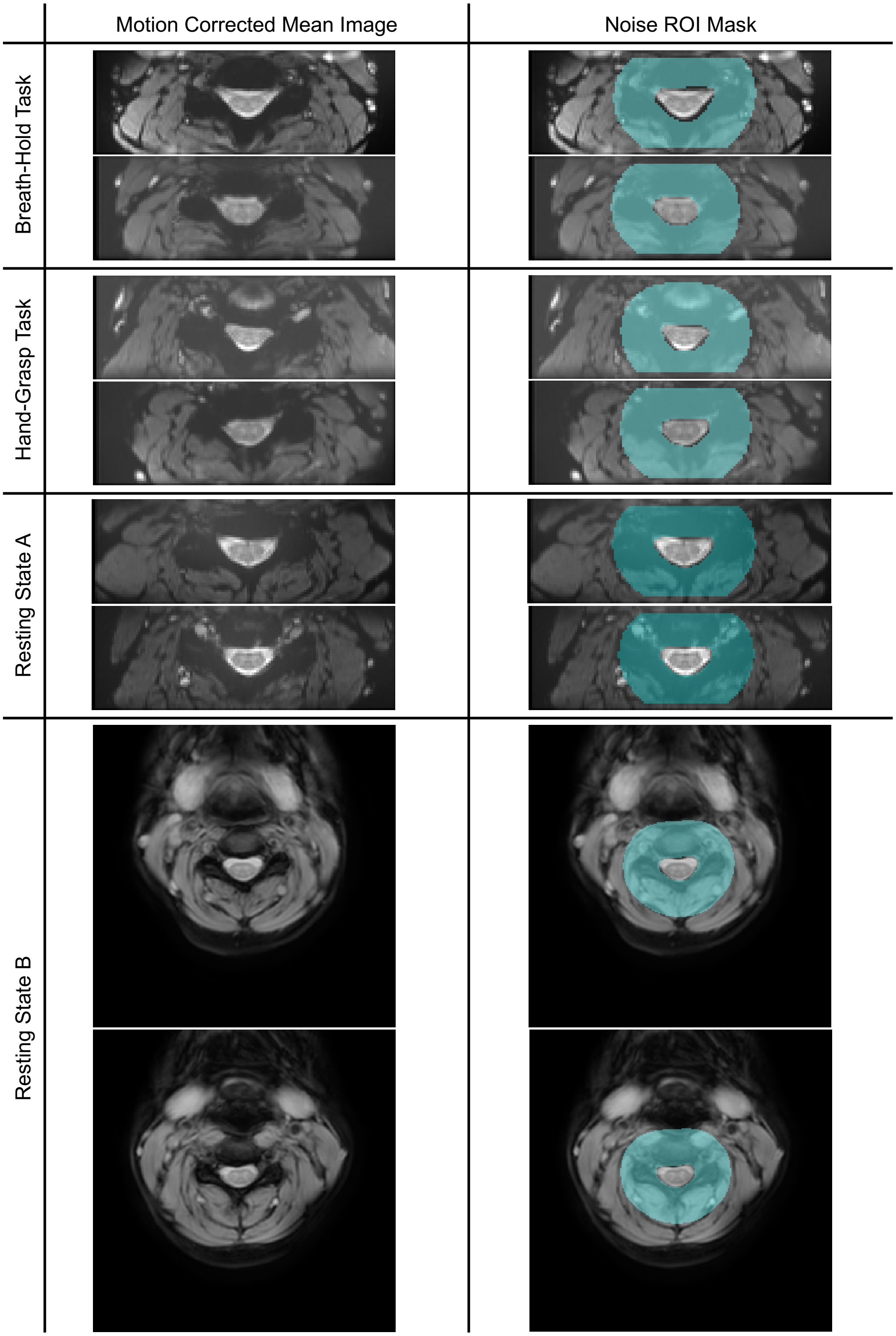


**Fig. S2. Example noise ROIs for the breath-hold task, hand-grasp task, and both resting state datasets.** The motion corrected mean functional image, and the noise ROI mask overlaid on the mean are shown for two example subjects per dataset. Breath-hold, hand-grasp, and resting state A fMRI data were collected with an inner field of view sequence.

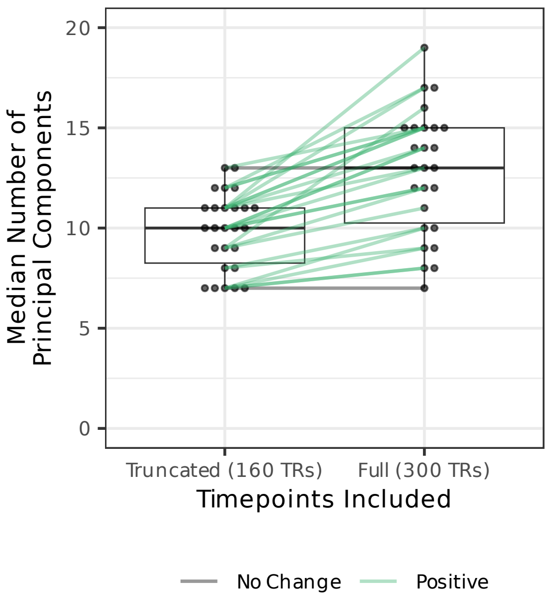


**Fig. S3. Comparison of median number of cutoff principal components for truncated and full hand-grasp task runs.** ‘Truncated’ vs. ‘Full’ represents the number of timepoints included in the parallel analysis. One run per subject is shown. Lines connect datapoints for the same subject/run and are colored by the direction of change.

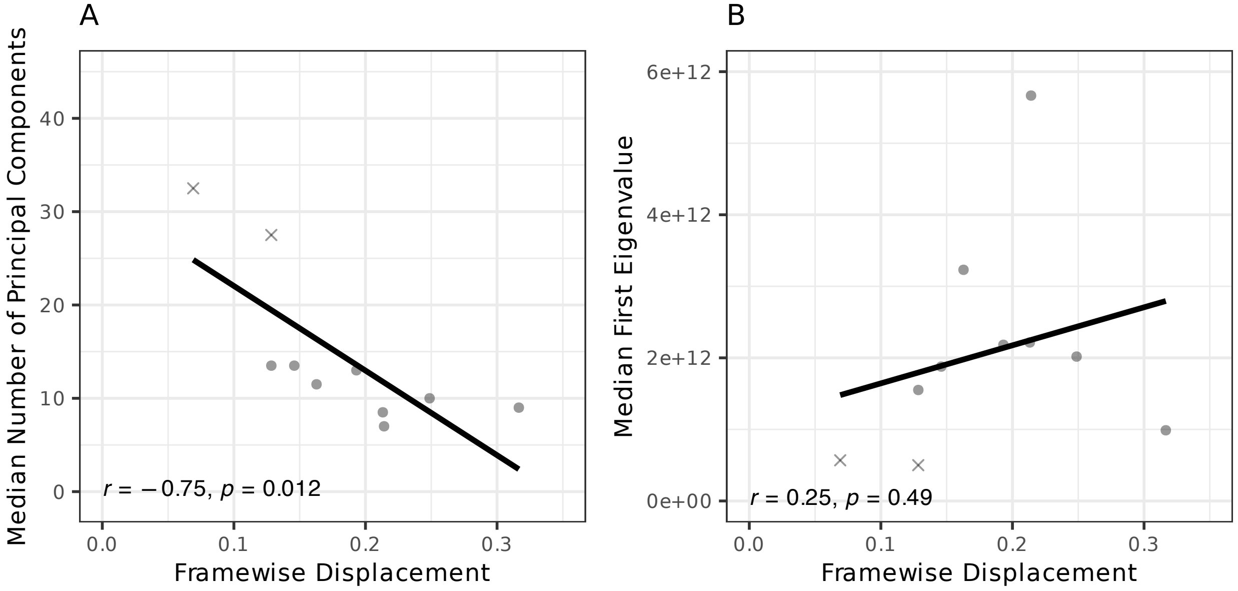


**Fig. S4. Comparison of resting state motion to the median number of cutoff principal components and the first eigenvalue. (A)** The in-plane motion (framewise displacement; X and Y motion) vs. median number of principal components for each subject’s run in the resting state B dataset. **(B)** The in-plane motion vs. the eigenvalue associated with the first principal component (median across slices). In both panels, the outliers identified in **Fig. 1** identified with an x.

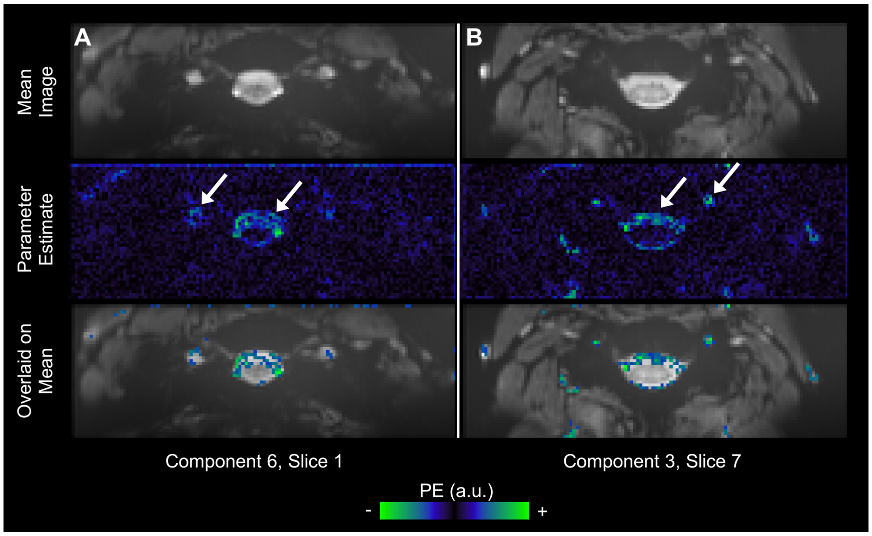


***Fig. S5. Parameter estimate fits for SpinalCompCor regressors in an example subject from the hand-grasp dataset (Extended Model).*** *The top and middle rows of panels A and B show the same Mean Image and Parameter Estimate as Fig. 3. The bottom row shows the parameter estimate with an arbitrary threshold overlaid on the mean image to illustrate the proximity of higher parameter estimates to the CSF and white matter.*

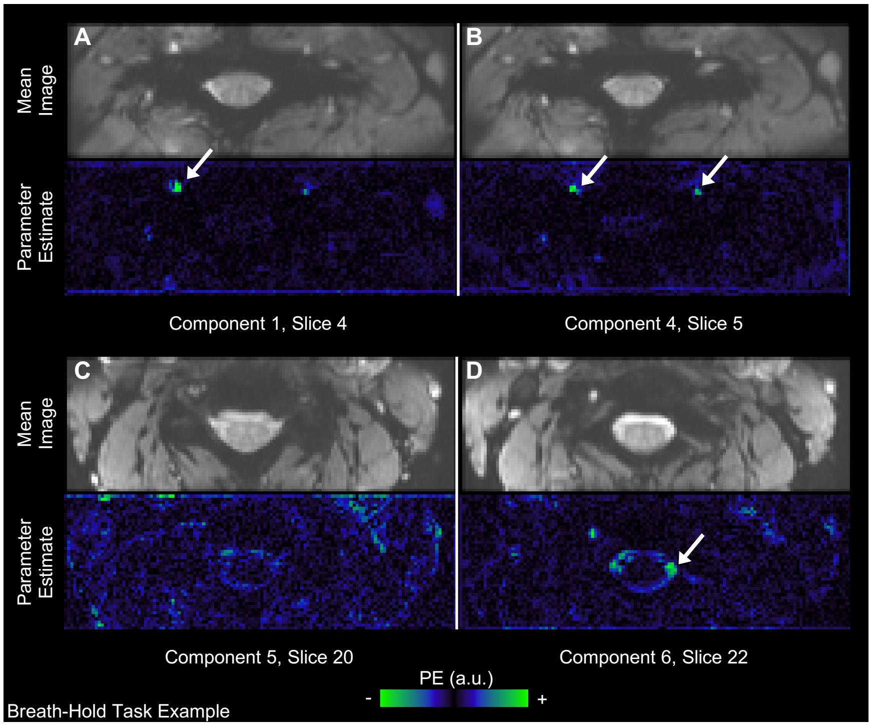


***Fig. S6. Parameter estimate fits for SpinalCompCor regressors in an example subject from the breath-hold task dataset.*** *The motion corrected mean image and corresponding principal component parameter estimate maps are shown for each example component/slice. Arrows point out areas with stronger parameter estimates. There are 25 slices in this acquisition, numbered 0-24. (PE=parameter estimate, a.u.=arbitrary units)*

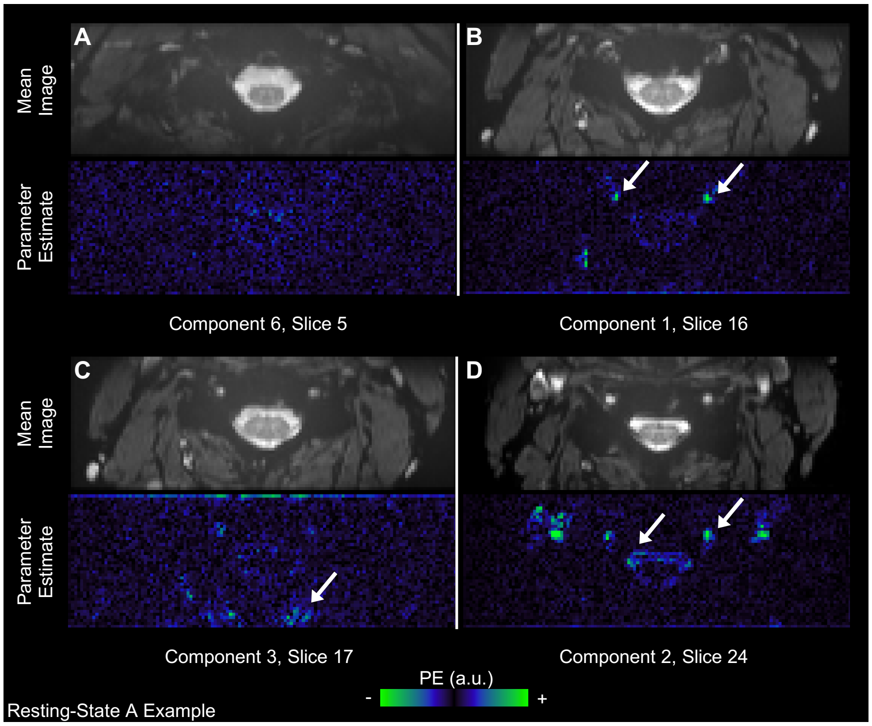


***Fig. S7. Parameter estimate fits for SpinalCompCor regressors in an example subject from the Resting State A dataset.*** *The motion corrected mean image and corresponding principal component parameter estimate maps are shown for each example component/slice. Arrows point out areas with stronger parameter estimates. There are 31 slices in this acquisition, numbered 0-30. (PE=parameter estimate, a.u.=arbitrary units)*

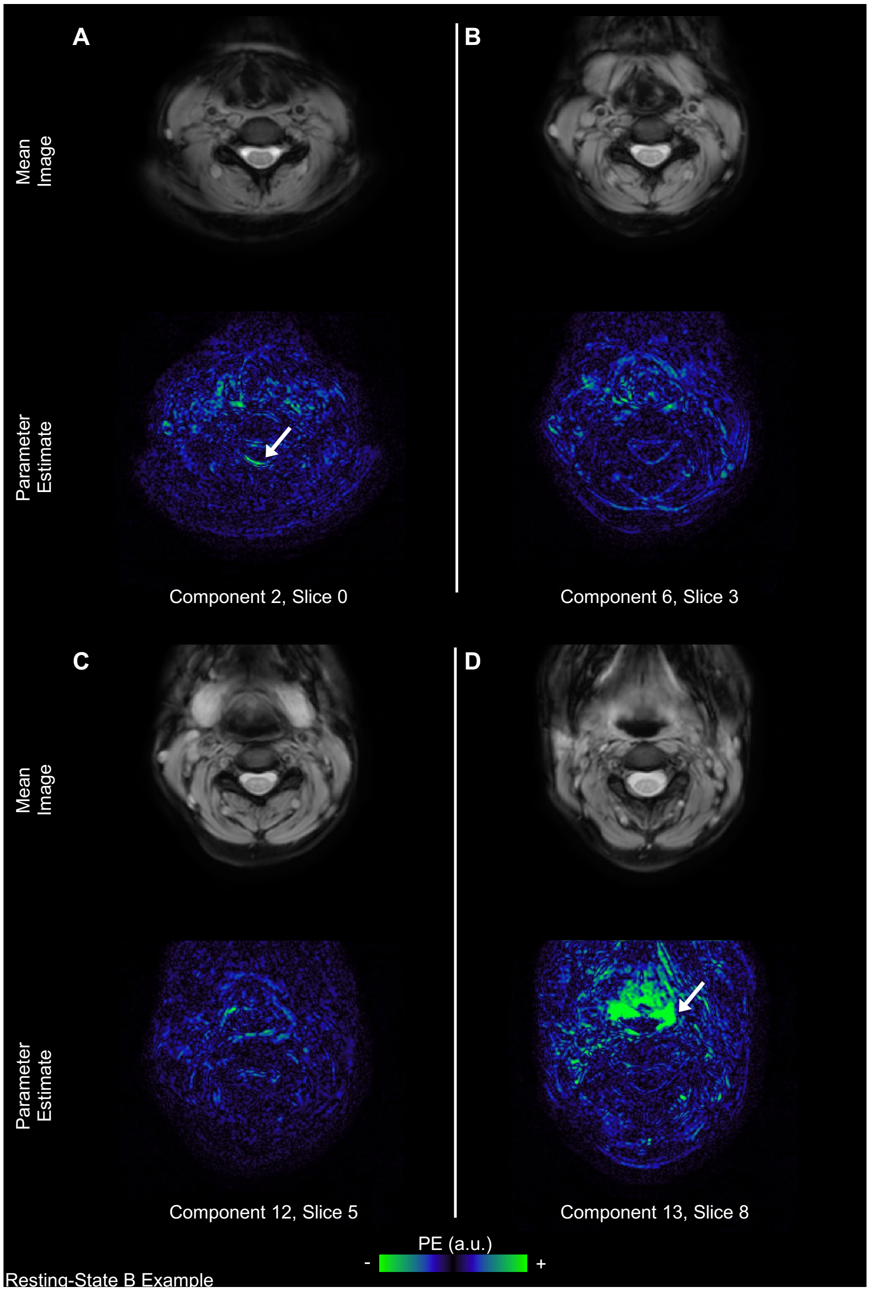


***Fig. S8. Parameter estimate fits for SpinalCompCor regressors in an example subject from the Resting State B dataset.*** *The motion corrected mean image and corresponding principal component parameter estimate maps are shown for each example component/slice. Arrows point out areas with stronger parameter estimates. There are 12 slices in this acquisition, numbered 0-11. (PE=parameter estimate, a.u.=arbitrary units)*

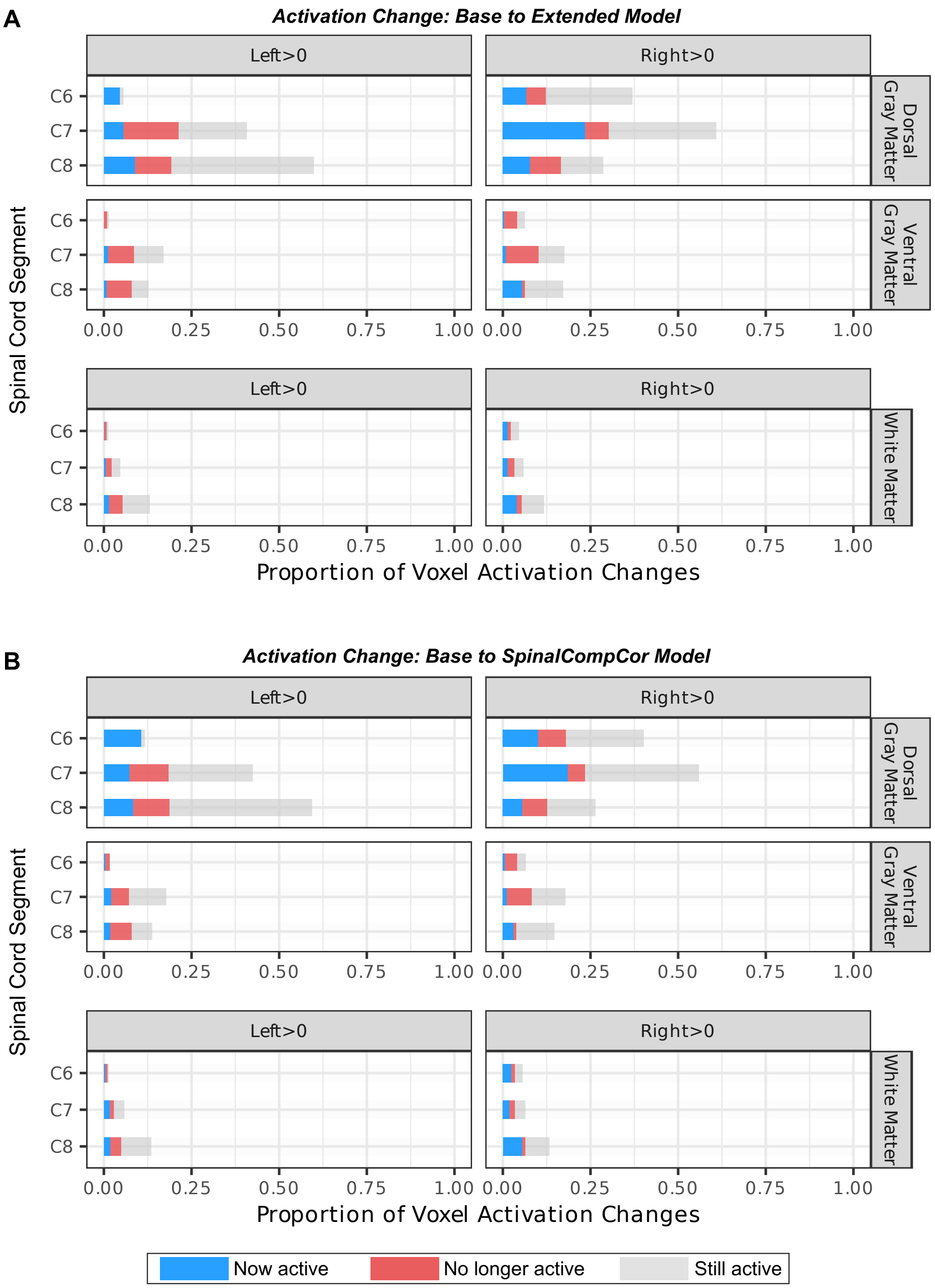


***Fig. S9. Voxel-wise activation changes for the hand-grasp task from the Base model in ROIs.*** *The proportion of voxels with the* ***(A)*** *Extended model or the* ***(B)*** *SpinalCompCor model that are now active (blue), are no longer active (red), or are still active (gray) are shown.*

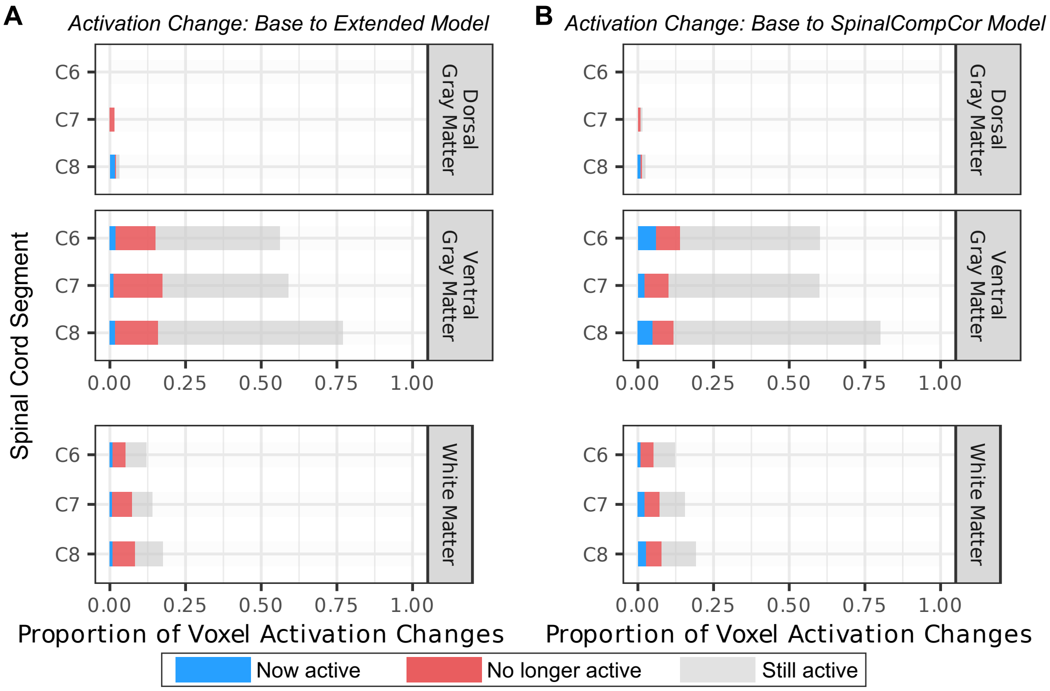


**Fig. S10.** ***Voxel-wise activation changes for the breath-hold task from the Base model in ROIs.*** *The proportion of voxels with the* ***(A)*** *Extended model or the* ***(B)*** *SpinalCompCor model that are now active (blue), are no longer active (red), or are still active (gray) are shown.*

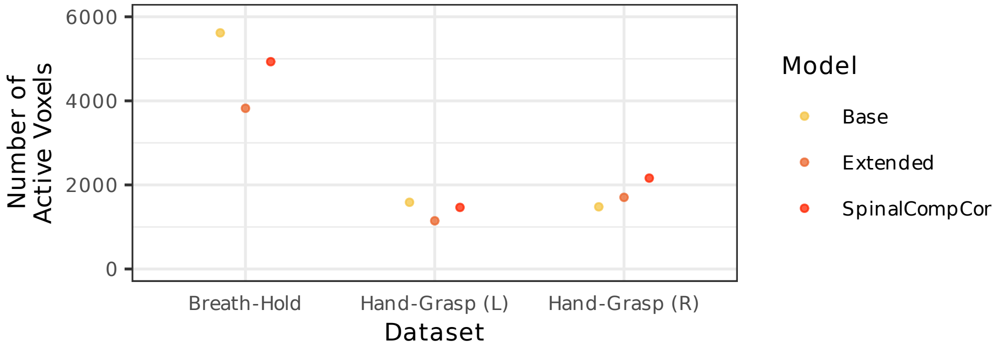


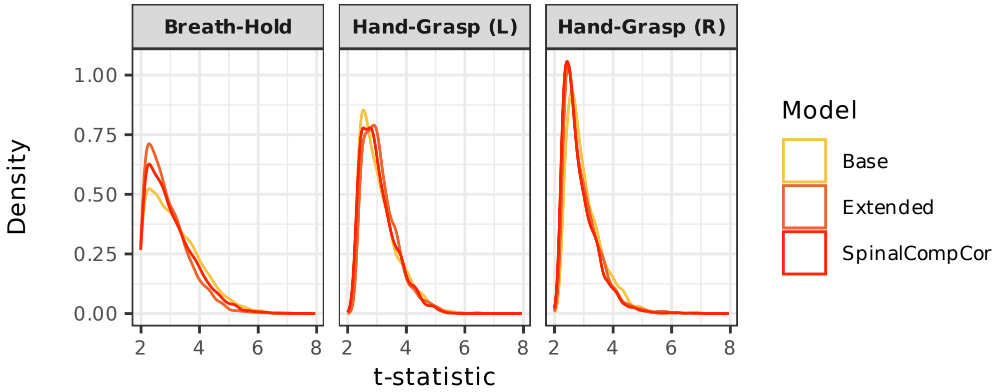


***Fig. S11. Significant voxels in breath-hold and hand-grasp group-level activation maps. (Top)*** *The number of significantly active voxels for each model.* ***(Bottom)*** *Density plot showing the distribution of t-statistics in significant voxels for each model.*

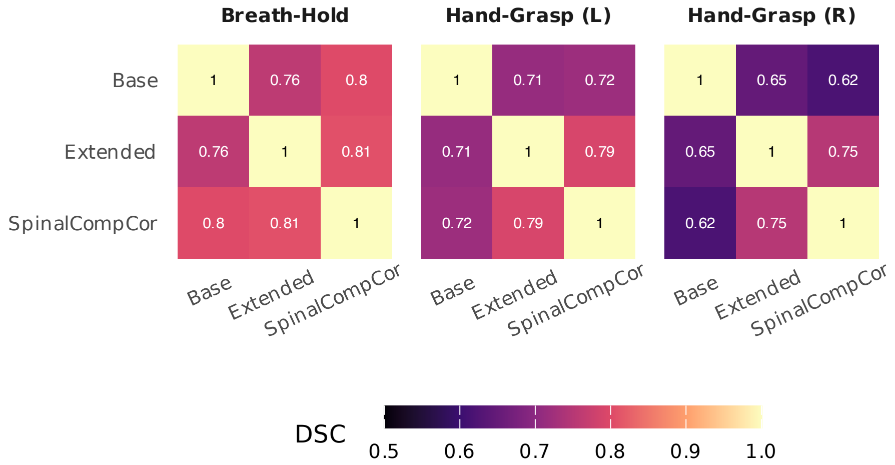


***Fig. S12. Spatial similarity of group-level activation between models.*** *Dice similarity coefficient (DSC) between models within masks of significantly active voxels.*

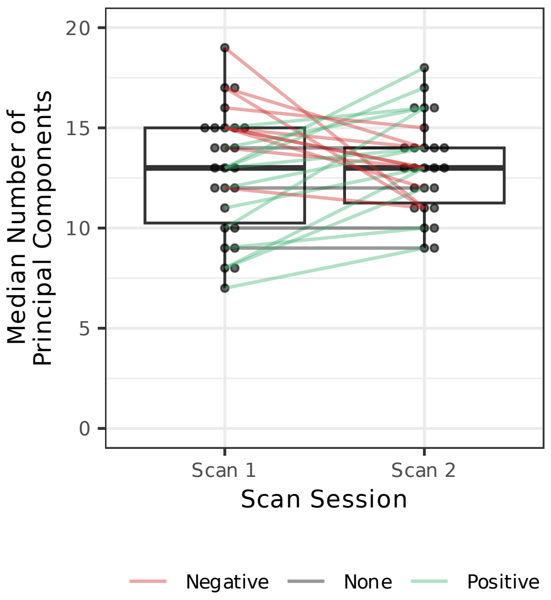


**Fig. S13. Comparison of median number of cutoff principal components for different runs (Scan 1 vs. Scan 2) for the hand-grasp task dataset.** Lines connect datapoints for the same subject and are colored by the direction of change.

**Table S1. Inclusion of regressors for noise modeling in recent human spinal cord fMRI studies.** Motion includes motion parameters from motion correction and/or motion outliers. CSF regressors were typically calculated as the top percent variance in a CSF mask, as the mean signal, or using PCA methods. Published studies in the past two years (from November 2, 2024) which detailed their noise and fMRI modeling methods were included. (RETROICOR=retrospective image correction (Glover et al., 2000), PNM=physiological noise modelling (Brooks et al., 2008), CSF=cerebrospinal fluid, RVT=respiration volume per time, HRV=heart rate variability).

| | | | **Included Regressors** | | | |
|---|---|---|---|---|---|---|
| **Authors (Year), Journal** | **Title** | **Task/Rest** | **RETROICOR / PNM** | **Motion** | **CSF** | **Other** |
| Dabbagh et al. (2024), *Imaging Neuroscience* | Reliability of task-based fMRI in the dorsal horn of the human spinal cord | Thermal Stimulus | X | X | X | |
| Seifert et al. (2024), *Hum Brain Mapp* | Thermal stimulus task fMRI in the cervical spinal cord at 7 Tesla | Thermal Stimulus | X | X | X | |
| Kowalczyk et al. (2024), *Hum Brain Mapp* | Spinal fMRI demonstrates segmental organisation of functionally connected networks in the cervical spinal cord: A test-retest reliability study | Resting state | X | X | X | |
| Braaß et al. (2023), *Hum Brain Mapp* | Association between activity in the ventral premotor cortex and spinal cord activation during force generation-A combined cortico-spinal fMRI study | Motor Task | X | X | X | RVT<br>HRV |
| Combes et al. (2023), *Sci Rep* | Detection of resting-state functional connectivity in the lumbar spinal cord with 3T MRI | Resting State | X | X | X (PCA) | Not spine ROI (PCA) |
| Hemmerling et al. (2023), *Hum Brain Mapp* | Spatial distribution of hand-grasp motor task activity in spinal cord functional magnetic resonance imaging | Motor Task | X | X | X | $P_{ET}CO_2$<br>SpinalCompCor |
| Kinany et al. (2023), *NeuroImage* | Decoding cerebro-spinal signatures of human behavior: Application to motor sequence learning | Motor Task | X | X | X | Mean global signal outside the spinal cord |
| Landelle et al. (2023), *Mov Disord* | Altered Spinal Cord Functional Connectivity Associated with Parkinson's Disease Progression | Resting State | X | X | X (PCA) | RVT<br>HRV |
